# Ultradeep characterisation of translational sequence determinants refutes rare-codon hypothesis and unveils quadruplet base pairing of initiator tRNA and transcript

**DOI:** 10.1101/2022.05.02.490318

**Authors:** Simon Höllerer, Markus Jeschek

## Abstract

Translation is a key determinant of gene expression and an important biotechnological engineering target. In bacteria, 5’-untranslated region (5’-UTR) and coding sequence (CDS) are well-known mRNA parts controlling translation and thus cellular protein levels. However, the complex interaction of 5’-UTR and CDS has so far only been studied for few sequences leading to non-generalisable and partly contradictory conclusions. Herein, we systematically assess the dynamic translation from over 1.2 million 5’-UTR-CDS pairs in *Escherichia coli* to investigate their collective effect using a new method for ultradeep sequence-function mapping. This allows us to disentangle and precisely quantify effects of known and hypothetical sequence determinants of translation. We find that 5’-UTR and CDS individually account for 53% and 20% of variance in translation, respectively, and show conclusively that, contrary to a common hypothesis, tRNA abundance does not explain expression changes between CDSs with different synonymous codons. Moreover, the obtained large-scale data clearly point to a base-pairing interaction between initiator tRNA and mRNA beyond the anticodon-codon interaction, an effect that is often masked for individual sequences and therefore inaccessible to low-throughput approaches. Our study highlights the indispensability of ultradeep sequence-function mapping to accurately determine the contribution of parts and phenomena involved in gene regulation.

## INTRODUCTION

Translation is a key step of gene expression and an important engineering target in synthetic biology. To this end, genetic parts that influence translation are modified to alter absolute and relative expression levels to engineer biosystems through control of individual genes, pathways and even entire metabolic networks (1–3). In prokaryotes, initiation of translation is the rate-limiting step in the translational process, during which ribosomes assemble on the mRNA to start the templated elongation of the nascent polypeptide (4–7). At the onset of this step, the 30S ribosomal subunit attaches to the ribosome binding site (RBS) in the 5’-untranslated region (5’-UTR) upstream of the coding sequence (CDS). The 3’-end of the 16S rRNA hybridises with the Shine-Dalgarno (SD) motif, a conserved five to eight nucleotide (nt) sequence located upstream of the start codon, which facilitates translation (8–12). However, since Shine and Dalgarno’s discovery in 1973 (10), various additional influencing factors and sequence determinants affecting translation initiation were identified. For example, the distance between SD motif and start codon, the type of start codon, and interactions between distant 5’-UTR parts and the ribosome play important roles (13–21). Remarkably, in some cases SD-like motifs are not required for translation, an observation hinting at the existence of other mechanisms besides “canonical” translation initiation (22–26). Further, the influence of mRNA secondary structures was studied under the hypothesis that the required unfolding of such structures during translation initiation might decrease expression (14,20,27–40). For example, stable secondary structures around the start codon were found to hinder translation, while structures further up- or downstream had less pronounced effects (35).

Moreover, codon usage was found to influence translation. Genome-wide analyses of *E. coli* and other organisms revealed an overrepresentation of rare codons in the first five to ten triplets of the CDS in native genes, and their occurrence in this region was found to coincide with high expression (29,41–44). These observations led to two different hypotheses that differ fundamentally in terms of the underlying causality. The first hypothesis is related to the fact that cellular tRNA concentrations correlate with the occurrence frequency of their cognate codons (45–47). It was postulated that rare codons (with low-abundant cognate tRNAs) may have been evolutionary selected for within the N-terminal CDS to slow down early translation elongation and reduce premature termination due to clashing ribosomes (38,48-52). These “translational ramps” were postulated to be causally responsible for elevated expression of genes rich in rare codons at the CDS’s 5’-end. As an alternative explanation independent of tRNA abundance, a second hypothesis has been proposed based on the fact that many rare codons are (or happen to be) AT-rich (29,52). Their occurrence is therefore associated with a lower tendency to form stable mRNA secondary structures (29,33,43), which are known to hinder translation initiation.

In the context of these two hypotheses, several studies have been conducted to investigate the impact of codon usage on expression focussing either on the N-terminal codons alone (29,33,37,43,44) or the entire CDS (29) while applying different metrics of codon usage such as the codon adaptation index (CAI) (41), the frequency of “optimal” codons pairing with the most abundant tRNAs (45), and the tRNA adaptation index (tAI) (53), as discussed in detail elsewhere (44,54-56). Remarkably, while there is clear evidence for a high degree of interactivity between 5’-UTR and CDS, these two mRNA parts were handled separately in these studies: commonly only one of the two parts (either 5’-UTR or CDS) was diversified at a time, and systematic testing of larger numbers of 5’-UTR-CDS combinations to assess their interaction was not performed (15,20,33). Thus, due to the strong interdependence the measured effects could not be clearly assigned to individual sequence parameters, and their contribution to overall expression could not be accurately quantified. Moreover, many early studies relied on experimental testing of only a few “hand-picked” sequences (usually less than 100 variants) due to limitations in experimental throughput or library generation (note that the CDS cannot be freely mutated, since of amino acid substitution may result in change or loss of reporter protein activity). Although valuable contributions, such empiric efforts have proven insufficient to establish generalisable rules and quantitative measurements for the potential effects of sequence parameters, which in some cases even led to contradictory conclusions. For example, the question of whether tRNA abundance has a significant impact on translation or whether the observed effect is caused by mRNA secondary structures alone remains inconclusively answered (29, 54). Enabled by advances in DNA synthesis and sequencing, some recent works assessed larger numbers of 5’-UTRs or CDSs, again only diversifying one of the two sequence parts at a time (20,33,36,38,57,58). In a recent study, Arkin and co-workers combined full-factorial *in silico* design with DNA synthesis on arrays to evaluate the principles of sequence design for translation in a systematic manner (37). They tested synthetic sequences combining a single bicistronic 5’-UTR (15,59) with 244,000 CDSs using fluorescence-activated cell sorting combined with next-generation sequencing (NGS). Several relevant sequence parameters such as AT-content, codon usage, and mRNA folding were varied and combined in a statistically full-factorial manner. This was achieved using a sophisticated modular design approach based on *a priori* hypotheses, which, however, bears the risk of introducing “user-borne” bias.

Herein, we describe our efforts to overcome the prevailing lack of knowledge about the impact of different mRNA parts and sequence parameters on translation with the goal to assess and accurately quantify their effect. We combine randomly generated 5’-UTRs and CDSs following different assembly strategies to obtain libraries of random, combinatorial and full-factorial 5’-UTR-CDS combinations. Using a recently developed method for ultradeep sequence-function mapping (58), we dynamically assess translation of more than 1.2 million 5’-UTR-CDS pairs in more than 8.8 million sequence-function data points and different genetic backgrounds. The extremely high throughput and the modular assembly strategy applied herein allow us to systematically disentangle and assess individual and combined effects of 5’-UTR and CDS, and to quantify the contribution of various sequence parameters including individual bases and positions, mRNA secondary structures, 16S-rRNA hybridisation, and codon usage.

## MATERIAL AND METHODS

### Reagents

All chemicals were obtained from Sigma Aldrich (Buchs, Switzerland). Restriction enzymes were obtained from New England Biolabs (Ipswich, USA). PCR was performed using Q5 DNA polymerase from New England Biolabs (Ipswich, USA). Oligonucleotides (**Suppl. Tab. 1**) were obtained from Microsynth AG (Balgach, Switzerland). All primers containing degenerate bases were ordered PAGE-purified. Custom duplex DNA adapters and gene fragments were obtained from Integrated DNA Technologies (Leuven, Belgium). Plasmid DNA for cloning was extracted with the ZR Plasmid Miniprep kit from Zymo research (Irvine, USA). Plasmid DNA from cultures used for subsequent sample preparation for NGS was extracted with the QIAprep Spin Miniprep kit from Qiagen (Hilden, Germany). Gel extraction of DNA was performed using Zymoclean Gel DNA Recovery Kits from Zymo research (Irvine, USA).

### Strains, cultivation conditions and growth analysis

*Escherichia coli* (*E. coli*) TOP10 *ΔrhaA* (l-rhamnose isomerase) was used throughout the study. The generation of these rhamnose utilisation-deficient strain is described elsewhere (58). For experiments with plasmid-borne variants of tRNA^fMet^, the strain *E. coli* TOP10 *ΔrhaA ΔmetZWV* was generated by additional replacement of the chromosomal *metZWV* locus with a spectinomycin resistance cassette using the method described by Datsenko and Wanner (60). The spectinomycin resistance cassette was PCR-amplified from a commercial gene fragment (**Suppl. Note 1**) using primers p1 and p2 (**Suppl. Tab. 1**) to generate the linear fragment for transformation complementary to 41 bp both up- and downstream of the chromosomal *metZWV* locus. Transformants were verified for successful integration by colony PCR using primers p3 and p4 and subsequent Sanger sequencing. The exact genotypes of both *E. coli* strains are provided in **Supplementary Table 2**. *E. coli* cells were generally cultivated in lysogeny broth (LB) supplemented with 50 mg L^-1^ kanamycin, 50 mg L^-1^ streptomycin and 10 g L^-1^ D-glucose for repression of the rhamnose-inducible promoter. 15 g L^-1^ agar were added for plate cultures. Cells were grown at 37 °C in an incubator (plates) or shaking incubator at 200 rpm (shake flasks cultivations). Doubling times of strains with different tRNA^fMet^ variants were determined in biological triplicate cultures as follows. *E. coli* TOP10 *ΔrhaA* (“WT”) and *E. coli* TOP10 *ΔrhaA ΔmetZWV (“ΔmetZWV”*) were transformed with pSEVA361 (empty vector), ptRNA^fMet-A37^, ptRNA^fMet-A37G^ or ptRNA^fMet-A37U^, respectively. Sequence-verified transformants of each strain were used to inoculate an overnight pre-culture in LB (34 mg L^-1^ chloramphenicol; 12.5 mg L^-1^ spectinomycin for *ΔmetZWV*). After, 120 mL main cultures in baffled shake flasks (1 L) were inoculated to a starting OD_600_ of 0.01 and incubated shaking (37 °C, 200 rpm). The OD_600_ was measured in intervals of 15-30 minutes and doubling times were determined by dividing ln(2) by the specific growth rate during exponential growth.

### Plasmid and library construction

A list of plasmids used in this study is provided in **Supplementary Table 3**. Plasmids were constructed by conventional restriction-ligation cloning. To enable facile library cloning, plasmid pASPIre4 (**Suppl. Fig. 1**) was generated as a derivative of the previously published pASPIre3 (58). pASPIre4 additionally contains a SpeI restriction site within the CDs of *bxb1* to enable diversification of the 5’-UTR and codons 2-16 of *bxb1*.

Library inserts were generated by PCR with degenerate primers to diversify the respective regions and inserted into the pASPIre4 backbone thereafter. The fully randomised 5’-UTR-CDS library was generated via PCR using pASPIre4 as template and primers p5 and p6. After, the PCR product and pASPIre4 were digested with SpeI and PstI (37 °C, 3 h), gel purified and ligated (16 °C, T4 ligase, overnight). The ligation mixture was purified and used to electroporate freshly prepared *E. coli* TOP10 *ΔrhaA* cells (61). After 60 min recovery at 37 °C in LB with 10 g L^-1^ D-glucose, transformants were plated in different dilutions for colony counting on LB agar plates (50 mg L^-1^ kanamycin, 50 mg L^-1^ streptomycin and 10 g L^-1^ D-glucose). After overnight incubation (37 °C), 10 mL LB were added to the plates and approximately 400,000 colonies were scraped off with a spatula. Glycerol was added to the cell suspension to a final concentration of 150 g L^-1^ and the optical density at 600 nm (OD_600_) of the glycerol stock was adjusted to 5.0 before freezing of aliquots in liquid nitrogen and storage at −80 °C. This pool of clones was designated Lib_random_ and the corresponding plasmid architecture was termed pASPIre4_lib_ (**Suppl. Fig. 2**). For the uASPIre with mutated tRNA^fMet^ variants, a glycerol stock of Lib_random_ was plated on LB agar and plasmid DNA of approximately 50,000 clones was extracted and subsequently used to transform *E. coli* bearing the respective plasmids for the expression of tRNA^fMet^ (see below).

Combinatorial and full factorial libraries combining different 5’-UTRs and CDSs were generated in a stepwise procedure as illustrated in **Supplementary Figure 3**. First, 5’-UTR and CDS half-libraries (**Suppl. Figs. 4, 5**) were cloned separately as described above. The 5’-UTR half-library was generated by PCR with primers p5 and p7 on pASPIre4 as template and subsequently inserted into the pASPIre4 backbone using PstI and NotI. Primer p7 introduces degeneracy in the 5’-UTR and a BbsI site between the randomised 5’-UTR and the NotI site (**Suppl. Fig. 3**). The CDS half-library was generated by PCR with primers p8 and p9 on pASPIre4 as template and inserted into the pASPIre4 backbone using PstI and NotI. Primer p8 introduces degeneracy in the CDS and a BbsI site between the CDS and the PstI site (**Suppl. Fig. 3**). Transformants of both half-libraries were plated separately in various dilutions. Depending on the libraries to be created afterwards, a desired number of colonies was scraped off with a spatula and plasmid DNA was extracted: for Lib_comb1_, approximately 1,000 colonies of the 5’-UTR half-library and approximately 1,000 colonies of the CDS half-library; for Lib_comb2_, approximately 100 colonies of the 5’-UTR half-library and approximately 10,000 colonies of the CDS half-library. For Lib_fact_, ten plates of approximately 100 colonies each of the 5’-UTR half-library and ten plates of approximately 100 colonies each of the CDS half-library were scraped off. In a second step, 5’-UTR and CDS half-libraries were combined to generate libraries Lib_comb1_, Lib_comb2_ and Lib_fact_. To achieve this, plasmid DNA from the different 5’-UTR half-libraries was PCR-amplified with primers p9 and p10 and the PCR product was digested with BbsI and PvuI. Subsequently, these half-libraries were ligated into plasmid backbones isolated from the individual CDS half-libraries via digestion with PvuI and BbsI. Note that the BbsI type IIS restriction site enables scarless joining of 5’-UTR and CDS half-libraries using ATGC (start codon ATG + first downstream base) as sticky ends for ligation. Lib_comb1_ (approx. 1,000 5’-UTRs combined with approx. 1,000 CDSs) and Lib_comb2_ (approx. 100 5’-UTRs combined with approx. 10,000 CDSs) were used to transform *E. coli* TOP10 *ΔrhaA* yielding approximately 1.5 million and 2.3 million colonies, respectively. Lib_fact_ was transformed in ten separate batches (ten times 100 5’-UTRs combined with 100 CDSs) yielding ten full-factorial sub-libraries. Each of these should contain a maximum of approximately 10,000 different 5’-UTR-CDS combinations, amongst which theoretically all 5’-UTRs are combined with all CDSs and *vice versa*. Colonies of these ten sub-libraries were scraped off plates and pooled to equivalent cell densities according to their OD_600_.

All plasmids for overexpression of tRNA^fMet^ variants are derivatives of pSEVA361 (62). We selected the chromosomal m*etY* locus including promoters and terminators of *E. coli* TOP10 as a scaffold since it is monocistronic and therefore simpler to mutate compared to the *metZWV* locus. In this scaffold we introduced an A-to-G point mutation at position 47 of the tRNA^fMet^ to match the sequence of *metZWV* (note that the *metY*-derived tRNA differs by this one base from *metZWV* tRNAs, which are three identical tRNA^fMet^ copies). The resulting monocistronic design was obtained as commercial gene fragment in four versions containing the wild-type base (A) as well as three mutants (C, G and T) at position 37 of tRNA^fMet^, respectively. The gene fragments were cloned into pSEVA361 (p15A replicon, chloramphenicol resistance) via KpnI and SpeI sites using standard procedures and sequence verified. The resulting plasmids were designated ptRNA^fMet-A37^, ptRNA^fMet-A37C^, ptRNA^fMet-A37G^ and ptRNA^fMet-A37U^ (**Suppl. Fig. 6**, **Suppl. Tab. 3**) and used to transform *E. coli* TOP10 *ΔrhaA* and *E. coli* TOP10 *ΔrhaA ΔmetZWV*. Note that transformants of ptRNA^fMet-A37C^ failed to grow and could thus not be included in further experiments. To assess the effect of tRNA^fMet^ mutations, *E. coli* TOP10 *ΔrhaA* and *E. coli* TOP10 *ΔrhaA ΔmetZWV* bearing the plasmids for tRNA overexpression were each co-transformed with the pool of 50,000 variants of Lib_random_ (see above).

### Library cultivation, sample preparation and NGS

The different libraries were separately grown in independent shake flask cultivations. Lib_fact_ was cultivated in two biological replicates. Cultivations were conducted in 600 mL LB with 50 mg L^-1^ kanamycin and, in case of tRNA^fMet^ overexpression, 34 mg L-1 chloramphenicol in 5 L baffled shake flasks. Pre-warmed (37 °C) LB was inoculated from glycerol stocks of the respective libraries to an initial OD_600_ of 0.05. Cultures were grown at 37 °C in a shaking incubator at 200 rpm. At an OD_600_ of approximately 0.5, expression of *bxb1* was induced by addition of 2 g L^-1^ l-rhamnose. Samples were drawn at 0, 95, 225, 290, 360 and 480 minutes after induction and immediately diluted in an excess of ice-cold PBS. Cell suspensions were centrifuged (4,000 g, 10 min, 4 °C) and pellets were snap frozen on dry ice. Afterwards, plasmid DNA was extracted and digested with SpeI and NcoI (4 h, 37 °C). Target fragments containing the 5’-UTR-CDS region and the Bxb1 recombination substrate were purified via gel electrophoresis (2.5% agarose). Afterwards, duplex DNA adapters for Illumina NGS with sample-specific indices (**Suppl. Tab. 4**) were ligated to the target fragments and full-length ligation products were purified via gel electrophoresis (2% MetaPhor agarose, Lonza, Basel, Switzerland). Purity and concentration of extracted fragments were determined using capillary electrophoresis (Fragment Analyser, Agilent) and samples were pooled in equimolar ratios. The pool was spiked with 15% PhiX DNA to increase sample diversity and afterwards sequenced on an Illumina NovaSeq6000 platform (SP flowcell, paired-end reading with at least 30 cycles forward and 100 cycles reverse read). Primary sequencing data were processed with Illumina RTA version V3.4.4 and bcl2fastq to obtain *.fastq files for further processing (see below).

### NGS data processing

NGS raw data analysis was performed using a combination of *bash* and *R* scripts (R version 4.1.2) running on a Red Hat Enterprise Linux Server (release 7.9). Annotated scripts for raw data processing will be made available upon final publication.

In brief, forward and reverse reads from *.fastq files were paired. From the forward reads, the identity of the sample-specific index (six options) and the state of the Bxb1 substrate (either unflipped or flipped), were extracted through alignment against all possible twelve combinations allowing a maximum of three mismatches between read and reference to avoid data loss due to sequencing errors. Afterwards, a similar procedure was applied to the reverse reads to identify the second sample-specific index (six options). Next, the sample-specific combination of forward and reverse indices was used to split the data and assigning reads to the different libraries and sampling time points (**Suppl. Tab. 5**).

Next, NGS reads with a frameshift within the CDS (e.g. due to sequencing errors or undesired mutations) were removed by filtering for the correct positioning of the constant first five nucleotides (ATGCG) of the *bxb1* CDS. Then, all 40 randomised nucleotides of 5’-UTR (25 nt) and CDS (each third nucleotide in codons 2-16; in total 15 nt) were extracted for each read, serving as unique identifier for each variant (i.e. 5’-UTR-CDS combination). To rescue reads with sequencing errors in the variable regions (less than 5% of total reads), a clustering procedure was applied to Lib_comb1_, Lib_comb2_ and Lib_fact_ to map them to actual (i.e. physically present) variants. This clustering can be applied since the extremely large theoretical sequence space of these variable regions (40 nt randomised; >10^23^ possible permutations) renders the occurrence of highly similar sequences virtually impossible. First, variants were sorted based on their total read number across all time points. Then, starting with the most frequent variant, all other variants with a Hamming distance of 1 (i.e. maximum of one substitution) were mapped back to this variant. This procedure was continued with the next most abundant variant until all remaining variants were further than one substitution apart from all others. 5’-UTRs and CDSs were treated separately to keep the computational complexity manageable. For Lib_random_, clustering was omitted since all 5’-UTRs and CDSs in this library are unique rendering the mapping process computationally infeasible. Afterwards, the number of reads with unflipped and flipped Bxb1 substrates was counted for the remaining variants and for each time sample to obtain time-resolved flipping profiles.

Lastly, an additional filtering step was performed to ensure high data quality, which excludes variants with less than 10 reads in at least one of the six time points. Moreover, variants containing an unintended non-synonymous codon mutation in the CDS were removed (227 variants).

This data processing procedure resulted in 1,214,438 high-quality variants split across the 4 libraries with an average of 464.3 reads per variant or 77.4 reads per variant and time point. For the uASPIre of tRNA^fMet^ mutants, this procedure resulted in 44,289 high-quality variants. In total, this amounts to 8,881,032 sequence-function pairs obtained from three NGS runs. The relative trapezoidal area under the flipping curve (termed “integral of the flipping profile”, IFP) was calculated for each variant. For Lib_fact_, the average IFP of the two biological replicates was used. Processed data and annotated scripts for data processing will be made available upon final publication.

### Correlation of Bxb1 recombination with cellular Bxb1-sfGFP levels

To convert Bxb1-catalysed flipping into relative cellular Bxb1 concentrations, we used the same approach as described previously, which relies on translational fusion of Bxb1 to the superfolder green fluorescent protein (sfGFP) and the use of internal standard RBSs (58). In brief, we first recorded the sfGFP fluorescence of 31 manually constructed RBSs controlling translation of the Bxb1-sfGFP fusion. These RBSs span a wide range of RBS strengths (from low to high) as previously shown in triplicate shake flask cultivations (58). A pool of these 31 standard RBSs was cultivated in a separate shake flask in parallel to the cultivations of Lib_random_, Lib_comb1_ and Lib_comb2_ and processed alongside the different libraries as described above. From the resulting NGS data, we obtained the IFP for the standard RBSs and constructed a calibration curve between IFP and the aforementioned sfGFP fluorescence measurements (58). A LOESS fit (locally estimated scatterplot smoothing) was used to correlate the IFP with the slope of the cell-specific sfGFP signal between 0 and 290 minutes after induction (slope GFP_0-290min_) using the function *loess* from the *R* package *stats*. Relying on the LOESS function, the IFP values of all library members were converted into the corresponding slope GFP_0-290min_. The resulting values were normalised to the maximum slope GFP_0-290min_ in the entire data and the normalised slope GFP_0-290min_ was designated relative translation rate (rTR) and used for all further analyses. Code and parameters of the LOESS fit will be made available upon final publication.

### Splitting of full-factorial sub-libraries

Since Lib_fact_ consists of ten full-factorial sub-libraries that were sequenced in bulk, the resulting data had to be computationally split into the sub-libraries for further analysis. Therefore, we sequenced at least three clones (reference variants) from each sub-library by Sanger sequencing covering both the randomised 5’-UTR and CDS regions. From the resulting reference sequences, we reconstructed and split the ten individual sub-libraries as follows: all variants that shared either the 5’-UTR or CDS with one of the reference sequences were assigned to the corresponding sub-library. To obtain full-factorial sub-libraries (i.e. libraries in which the majority of 5’-UTRs is combined with each CDS and vice versa), we further removed all variants with a 5’-UTR that occurred in combination with less than 50 CDSs as well as all variants with a CDS that occurred in combination with less than 50 5’-UTRs.

### Data analyses

Data analysis was conducted in *R* (version 4.1.2) and figures were produced using the package *ggplot2*. Scripts will be made available upon final publication.

For ANOVA of positional effects, variants from Lib_random_ were split according to their respective base in each of the 40 randomised positions within 5’-UTR and CDS (i.e. 40 splits for 40 position). After, type II ANOVA was performed using the R function *Anova* (package *car*) treating each positional group as covariate to determine the contribution of each covariate/position to the variance of the rTR in the entire library assuming additive behaviour. For the assessment of effects of single bases, we calculated the average rTR of all variants in Lib_random_ with a given base at a given position and divided the resulting value by the average rTR of all variants with any other base at this position. For example, the effect of U at 5’-UTR position −1 was calculated by dividing the average rTR of all variants with U at 5’-UTR position −1 (0.185) by the average rTR of all other variants (0.150). The resulting value (example: 1.233) represents the average relative in- or decrease in rTR for a given base and position. In the example above this means that the rTR of variants with U at 5’-UTR position −1 is on average 23.3% increased over the rest of the library. To assess the enrichment of bases amongst strong variants, variants in Lib_random_ were first split into two groups with rTR ≥ 0.5 (strong) and rTR < 0.5 (weak). After, the relative occurrence of each base at each position was calculated within each group. The ratio between the occurrences in the two groups represents the relative enrichment/depletion of a given base in a given position amongst strong variants over weak variants.

For calculations related to mRNA folding, bash scripts were used. Minimum free energy (mfe), ensemble free energy (efe) and mRNA accessibility (acc) were each calculated using two models for base pairing, the turner energy model (T) and the CONTRAfold model (C) (65,66), resulting in six different metrics (mfeT, mfeC, efeT, efeC, accT, accC). For mfeT and efeT, *RNAfold* (*ViennaRNA* package, version 2.4.18) and default parameters were used (67). For mfeC and efeC, and default parameters were applied. For accT and accC, the *Raccess* program was used (68). Next, Spearman’s correlation was calculated between each metric and the rTR. Note that Spearman’s correlation was used since rTR values do not follow a normal distribution (p-value of 1.11 × 10^-79^ according to Shapiro-Wilk normality test). Squared Spearman’s coefficient (ρ^2^) is reported as a measure of correlation between the respective folding metric and the ranked the rTR. Accordingly, the higher ρ^2^ of a metric, the more it explains the observed variance in the rTR. To identify the optimal mRNA sequence window that leads to the highest correlation between folding and rTR, mfeT and efeT were calculated for all possible sequence windows of lengths between 10 and 200 nucleotides within the first 200 positions of the mRNA. For computational reasons, this analysis was performed only on the 10’000 variants of Lib_random_ with the highest number of NGS reads. The best correlation between folding energy and rTR was achieved using the first 80 positions of the mRNA (i.e. between positions −27 and +53) (**Suppl. Fig. 7**). This “optimal” sequence window was then used to calculate mfeT, mfeC, efeT, efeC, accT and accC for all variants in all libraries. For accT and accC, the access length was set to 80 nucleotides in *Raccess*. For accessibility scanning, the correlation between the accessibility of each position and the rTR was determined applying an access length of 10 nucleotides in Raccess (accT_10nt_ and accC_10nt_).

To calculate 16S rRNA hybridisation energies, *RNAduplex* from the *ViennaRNA* package (67) was used, which only allows intermolecular base pairing. Allowing intramolecular base pairing would favour 5’-UTR-internal folds and thus disregard interactions with the 16S rRNA. Specifically, hybridisation energy was calculated between 5’-UTR (positional window: −18 to −4) and the 16S rRNA 3’-end (5’-ACCUCCUUA-3’). As an alternative, we also calculated a positional hybridisation energy between 16S rRNA 3’-end and a 9-nt sliding window along the entire mRNA.

The minimum edit distance was determined using the *stringdist* function of the *R* package *stringdist* and corresponds to the Levenshtein distance between the 7-bp long canonical SD motif AGGAGGU and a sliding 7-nt window within 5’-UTR positions −18 and −4. Levenshtein distance is the minimum number of operations (substitutions, deletions, and insertions) to transform one string into another.

The random forest model was built using *h2o.randomForest* from the R package *h2o* (https://github.com/h2oai/h2o-3). Variants of Lib_random_ were split into a randomly selected training set (90%) and a test set (10%), which was strictly held out during training. Sequences were encoded using one-hot encoding, a position-wise accessibility score accC_1nt_ (compare above), GC-content, minimum edit distance to the SD motif AGGAGGU, 16S rRNA hybridisation energy, the position of 16S rRNA hybridisation on the mRNA, as well as the folding metrics mfeT, mfeC, efeT, efeC, accT and accC (see above). Using tenfold cross-validation, the model was then trained with default parameters using 50 trees, and its performance was validated on the strictly held-out test set.

To quantify the contributions of UTR and CDS, we first grouped variants from Lib_comb1_, Lib_comb2_ and Lib_fact_ by their 5’-UTR and then calculated the average rTR of all CDSs in each group (i.e. rTR_UTR_). Similarly, we also grouped variants by their CDS and calculated the average rTR of all 5’-UTRs in each group (i.e. rTR_CDS_).

Codon adaptation index (CAI) and tRNA adaptation index (tAI) were calculated using the *cai* function from the *R* package *seqinr*. Codon weights and frequencies (**Suppl. Tab. 6**) were used as presented in Sharp *et al*. (41) and dos Reis *et al.*, respectively (53).

All sequence variants and their calculated parameters were combined into a single dataset and further analysed. This data set will be made available upon final publication.

### Data availability

Time series data including IFP and cellular Bxb1-sfGFP values (rTR, see above) for each variant including annotated scripts for data processing, statistical analyses and plotting will be made available upon final publication.

## RESULTS

### High-throughput characterisation of 5’-UTR-CDS combinations

It is challenging to investigate the impact of different mRNA parts on translation due to the vast sequence space of possible variants. For instance, even for a comparably short 5’-UTR of twelve nucleotides, more than 16 million (4^12^) sequences are possible. The sequence space becomes even larger if different parts are diversified simultaneously, which is required to analyse interactions and combined effects. Such combinatorial complexity cannot be addressed appropriately by measuring the expression of a few handpicked sequences. Instead, it requires high-throughput methodology capable of linking sequences to corresponding expression levels at large scale. To achieve this for 5’-UTR-CDS combinations, we capitalise herein on a recently developed technology for <u>u</u>ltradeep <u>A</u>cquisition of <u>S</u>equence-<u>P</u>henotype <u>I</u>nter<u>re</u>lations (uASPIre) (58). Briefly, uASPIre uses the phage recombinase Bxb1 to record functional information in DNA. This DNA-recorder enables, for instance, to determine both sequence and corresponding gene expression of gene regulatory elements via NGS at extremely high throughputs, which we have recently demonstrated in a proof-of-concept study (58).

To make uASPIre amenable for the characterisation of 5’-UTR-CDS combinations, we created the plasmid architecture shown in **Figure 1a**, which contains a gene encoding a Bxb1-sfGFP fusion (58) controlled by an l-rhamnose-inducible promoter (*P_rha_*) and a 150-bp stretch of silent DNA flanked by Bxb1’s cognate attachment sites *attB* and *attP* in opposite orientation (62). Furthermore, a SpeI site is introduced in codons 17 and 18 of the *bxb1* CDS via silent mutation (**Fig. 1a/b**), which enables facile exchange of the 5’-UTR and the first 16 codons of the *bxb1* CDS as well as NGS sample preparation (**Methods**). Once expressed, Bxb1-sfGFP converts its *attB*-/*P*-flanked DNA substrate from its initial (“unflipped” hereafter) to an inverted (“flipped” hereafter) state (**Fig. 1a**). Thus, Bxb1-sfGFP expression can be read out by determining the state of the substrate DNA by sequencing. Importantly, the flipping rate directly correlates with the cellular Bxb1-sfGFP concentration, and sequencing of many copies of this architecture via NGS can be used to determine the fraction of flipped DNA substrates (“fraction flipped” hereafter) amongst all copies of a given variant. This “oversampling” facilitates a precise, quantitative readout for Bxb1-sfGFP expression, whose resolution solely depends on the sequencing depth (i.e. number of reads obtained per variant) as we have previously shown (58).

**Figure 1:**
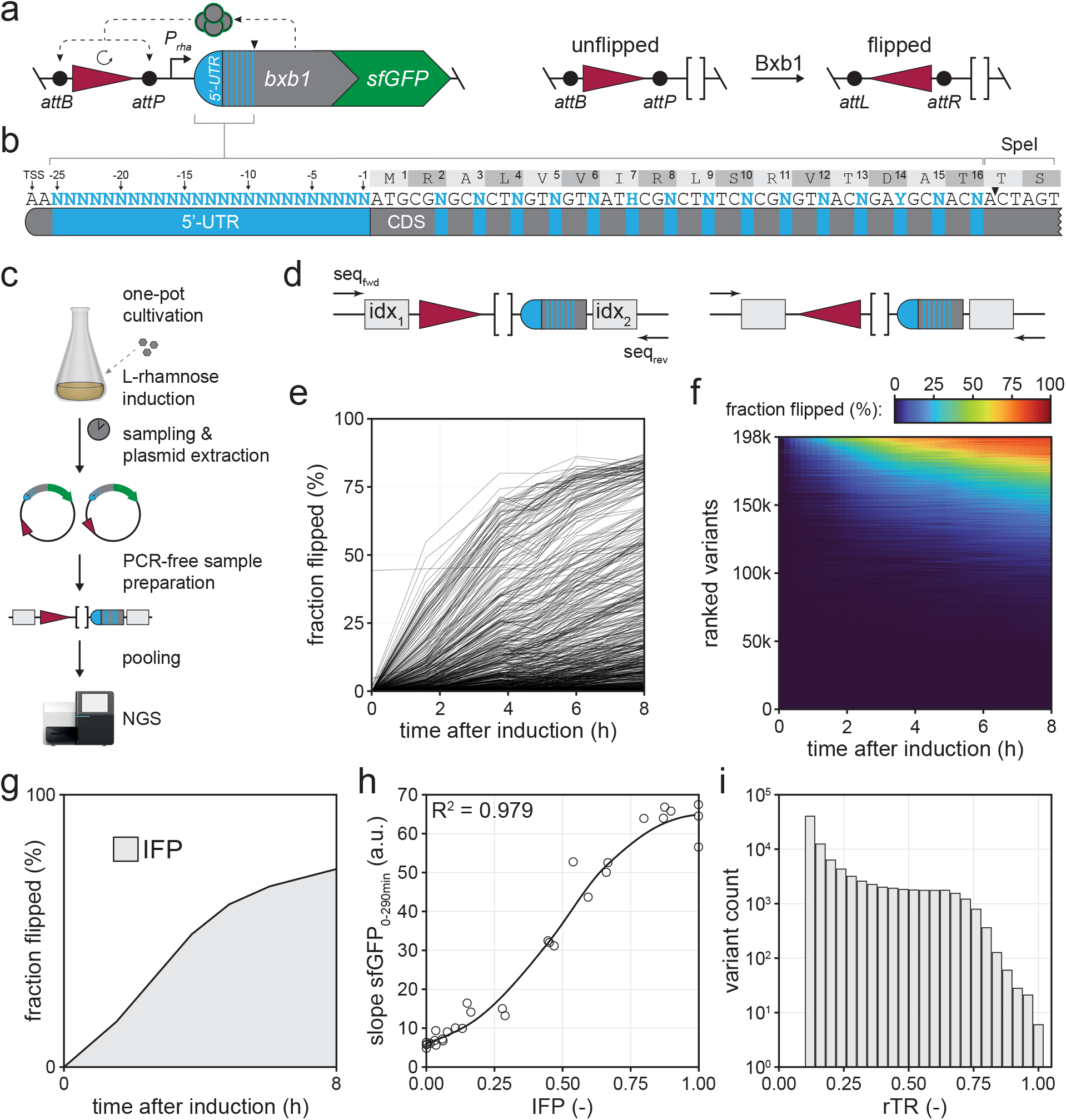
Ultradeep characterisation of 5’-UTR-CDS combinations. **a**) Plasmid architecture for the uASPIre of 5’-UTR-CDS pairs. A *bxb1-sfGFP* gene (translational fusion) controlled by *P_rha_* is placed on the same DNA molecule as the substrate modifiable by Bxb1-sfGFP, which is flanked by Bxb1 attachment sites (*attB/P*). A SpeI site in codons 17 and 18 of *bxb1-sfGFP* allows for seamless exchange of 5’-UTR and N-terminal CDS. Once expressed, Bxb1-sfGFP inverts its substrate from an unflipped into a flipped state creating recombined attachment sites (*attL/R*). **b**) Design of Lib_random_. The 25 nucleotides preceding the start codon are fully randomised. Additionally, the third positions of codons 2-16 are mutated allowing only synonymous codon replacements. Sequences follow the IUPAC nucleotide code (N: A/C/G/T, H: A/C/T, Y: C/T). TSS: transcriptional start site of *P_rha_*. **c**) Experimental workflow for the uASPIre of 5’-UTR-CDS pairs. Pooled transformants of Lib_random_ are grown in LB and *bxb1-sfGFP* expression is induced by l-rhamnose addition. After, samples are taken at different time points followed by plasmid extraction and preparation of NGS fragments followed by pooling of samples and NGS (**Methods**). NGS fragments are flanked by duplex adapters with sample-specific index combinations (grey boxes). **d**) Close-up view of target fragments for paired-end NGS using forward (seq_fwd_) and reverse (seq_rev_) sequencing primers. Forward reads are used to identify the first index (idx_1_) and the state of the recombinase substrate. Reverse reads are used to obtain the second index (idx_2_) and the sequence of 5’-UTR and CDS. **e**) Representative flipping profiles of 5’-UTR-CDS variants from Lib_random_. For clarity, only the 1,000 most abundant variants are displayed. **f**) Flipping profiles of all 198,174 Lib_random_ members above high-quality read-count threshold (**Methods**). Horizontal lines are time series of individual variants coloured according to the fraction flipped and ranked by the average fraction flipped across all time points from high (top) to low (bottom). **g**) Illustration of the IFP (grey area), i.e. the normalised trapezoidal integral of the flipping profile. **h**) Correlation between IFP and slope sfGFP_0-290min_ as shown for 31 standard RBSs (**Methods**). A LOESS function (black line) can be used to interconvert IFP and slope sfGFP_0-290min_ with high confidence. **i**) Histogram of the rTR of all variants from Lib_random_.

Next, we generated a first library through simultaneous diversification of the 5’-UTR and CDS of *bxb1-sfGFP* with the goal to characterise the impact on bacterial translation in a highly parallelised fashion relying on uASPIre (**Fig. 1b**, **Methods**). We mutated the 25 nucleotides directly upstream of the start codon applying full randomisation (i.e. N_25_-mer, N: equimolar mixture of A, C, G and T). This corresponds to the entire 5’-UTR in our setup except for two consecutive A’s at the 5’-end of the mRNA, which were fixed to match the native transcriptional start of *P_rha_* and thus avoid changes in transcription rates (69). Further, we mutated the third positions of codons 2-16 downstream of the start codon (ATG itself was kept constant) to additionally diversify the CDS. We selected this region since the first 30-50 nucleotides of CDSs reportedly affect translation whereas sequence changes further downstream show only negligible effects on expression (29,33). Importantly, in this region we only allowed synonymous (“silent”) codon replacements to maintain the same Bxb1 amino acid sequence and hence specific recombination activity for all library members, which is crucial to study only translational effects. This library is designated Lib_random_ hereafter pointing to the full randomisation of 5’-UTR and N-terminal CDS. Lib_random_ was used to transform *E. coli* yielding approximately 400,000 individual transformants. Specifically, we used the rhamnose-utilisation deficient strain TOP10 *ΔrhaA* to ensure temporally stable induction due to the lack of inducer consumption (58). Afterwards, transformants were pooled and cultivated in a single shake flask (**Fig. 1c**). In parallel, we cultivated 31 5’-UTR variants (“standard RBSs” hereafter) controlling the same *bxb1-sfGFP* fusion, which were constructed and characterised in a previous study (58). These standard RBSs span a wide range of expression levels and serve as internal standard sequences to compare different experiments. Further, they are used to convert the fraction flipped time series into practically more relevant metrics for protein expression relying on calibration curves generated from individual sfGFP fluorescence measurements (see below, **Methods**) (58). After induction by addition of l-rhamnose, six samples each were drawn over the course of eight hours from both cultures (Lib_random_ and standard RBSs), and plasmid DNA was extracted followed by NGS sample preparation (**Methods**). Note that sample preparation was carried without PCR amplification, which avoids non-linear PCR bias (58). The final target DNA fragments are flanked by NGS adapters with sample-specific indices and contain the DNA substrate modifiable by Bxb1 and the randomised 5’-UTR-CDS region. NGS adapters, substrate and 5’-UTR-CDS region were sequenced in an Illumina platform yielding approximately 10^8^ paired-end reads for Lib_random_. (**Fig. 1d**)

Next, we processed the NGS data to obtain time series of Bxb1-mediated flipping (“flipping profiles”) using a previously developed computational pipeline adapted to the new plasmid architecture (**Methods**)(58). This procedure yielded flipping profiles for 198,174 5’-UTR-CDS pairs above an applied minimal threshold of ten reads per time point and variant (i.e. high-quality data, average of 433.7 reads per variant). The base composition in Lib_random_ was homogeneously distributed across all diversified positions (**Suppl. Fig. 8**). Library members showed a diverse range of translational activities from low to high and a skew towards weaker variants as to be expected for full randomisation of the 5’-UTR (**Fig. 1e, f**) (70). Notably, the behaviour of the standard RBSs correlated strongly with results from our previous study even though the experiments were carried out approximately two years apart from each other (**Suppl. Fig. 9**)(58). This confirms the validity of the recorded data and indicates a high reproducibility and robustness of the uASPIre method in general. Next, we calculated the trapezoid integral of the flipping profiles (IFP, **Fig. 1g**), which constitutes a robust metric correlating well with rates of cellular Bxb1-sfGFP accumulation as previously shown (58). Indeed, the IFP of the 31 standard RBSs as determined in this study correlated well with the linear slope of the cell-specific Bxb1-sfGFP fluorescence between 0 and 290 minutes after induction (slope sfGFP_0-290min_, **Fig. 1h**, **Methods**). Therefore, IFP values can be converted into the slope sfGFP_0-290min_ relying on a fit applied between the two metrics for the standard RBSs. Specifically, we performed locally estimated scatterplot smoothing (LOESS) (**Fig. 1h**), and used the resulting fit function to convert the IFPs of Lib_random_ members into the corresponding slope sfGFP_0-290min_ normalised to the strongest variant found in this study (**Fig. 1i**, **Methods**). This normalised parameter was designated relative translation rate (rTR) and used for all further analyses, because it represents a practically more relevant metric for translational activity directly corresponding to cell-specific protein accumulation.

### Analysis of positional and base-specific effects on translation

Relying on the data generated for Lib_random_, we investigated the impact of different positions, nucleotides, and sequence motifs on expression. To assess positional effects, we performed analysis of variance (ANOVA) treating each variable position in the 5’-UTR (−25 to −1) and CDS (third positions of codons 2-16) as a covariate and calculated the contribution to the observed variance in rTR (**Fig. 2a, Methods**). Individual positions in the 5’-UTR explain between 0.3 and 1.5% of the variance. The most pronounced effect was observable for positions −13 to −8, which corresponds to an anticipated SD region, and, more unexpectedly, position −1. Within the CDS, the impact of codons decreases with increasing distance from the start codon with codon 2 showing the highest contribution (2.1%). Codons 2 to 8 show a marked effect, which strongly decreases to a negligible degree thereafter. Notably, the cumulative contribution of all 40 randomised positions only amounts to about 25% of which about 17.5% and 7.4% are attributed to 5’-UTR and CDS, respectively (**Suppl. Fig. 10**). The remaining high fraction of unexplained variance (about 75%) points towards a strong interaction between positions leading to non-additive behaviour. Next, we calculated the effect of specific bases at the variable positions by dividing the average rTR of variants with a given base at a position by the average rTR of all other variants (**Fig. 2b**). Generally, C and G tend to have a negative, and A and U a positive effect on translation, which is stronger in the 5’-UTR and weaker in the CDS decreasing with increasing distance to the start codon. A striking exception to that end are positions −14 to −7 (SD region), for which the effect of G is highly positive. The strongest negative effect is observable for CG<u>G</u> as the 2^nd^ codon (Arg) with corresponding variants being on average 26.3% weaker than those with CG<u>A</u>, CG<u>C</u> or CG<u>U</u> in this codon. The strongest positive impact is associated with U at 5’-UTR position −1 amounting to a mean rTR increase of 23.3%. Finally, to identify characteristic sequence determinants in particular of strong variants, we split the data from Lib_random_ into two sets of strong variants (i.e. rTR ≥ 0.5; 11,212 sequences) and weaker variants (i.e. rTR < 0.5; 186,962 sequences) and calculated the relative enrichment or depletion of each base at each position in the strong over the weaker subset (**Fig. 2c, Methods**). This analysis confirmed that both 5’-UTR and CDS of strong variants are generally enriched for A and U, and depleted for G and C except for a G-favouring region at positions −14 to −7. The latter shows a strong resemblance to archetypal AG-rich SD motifs, which commonly follow a consensus of AGGAGa/g in *E. coli*. In the CDS, we again observed a consistent decrease in positional importance with increasing codon number and a sharp drop of effect size after codon 8. Moreover, the aforementioned significance of U (but not A!) at 5’-UTR position -1 and the strong negative impact of CG<u>G</u> in codon 2 are confirmed by this analysis of strong sequences. The high and base-specific impact of these two positions prompted us to perform further analyses and experiments towards the causality of these effects (see below).

**Figure 2:**
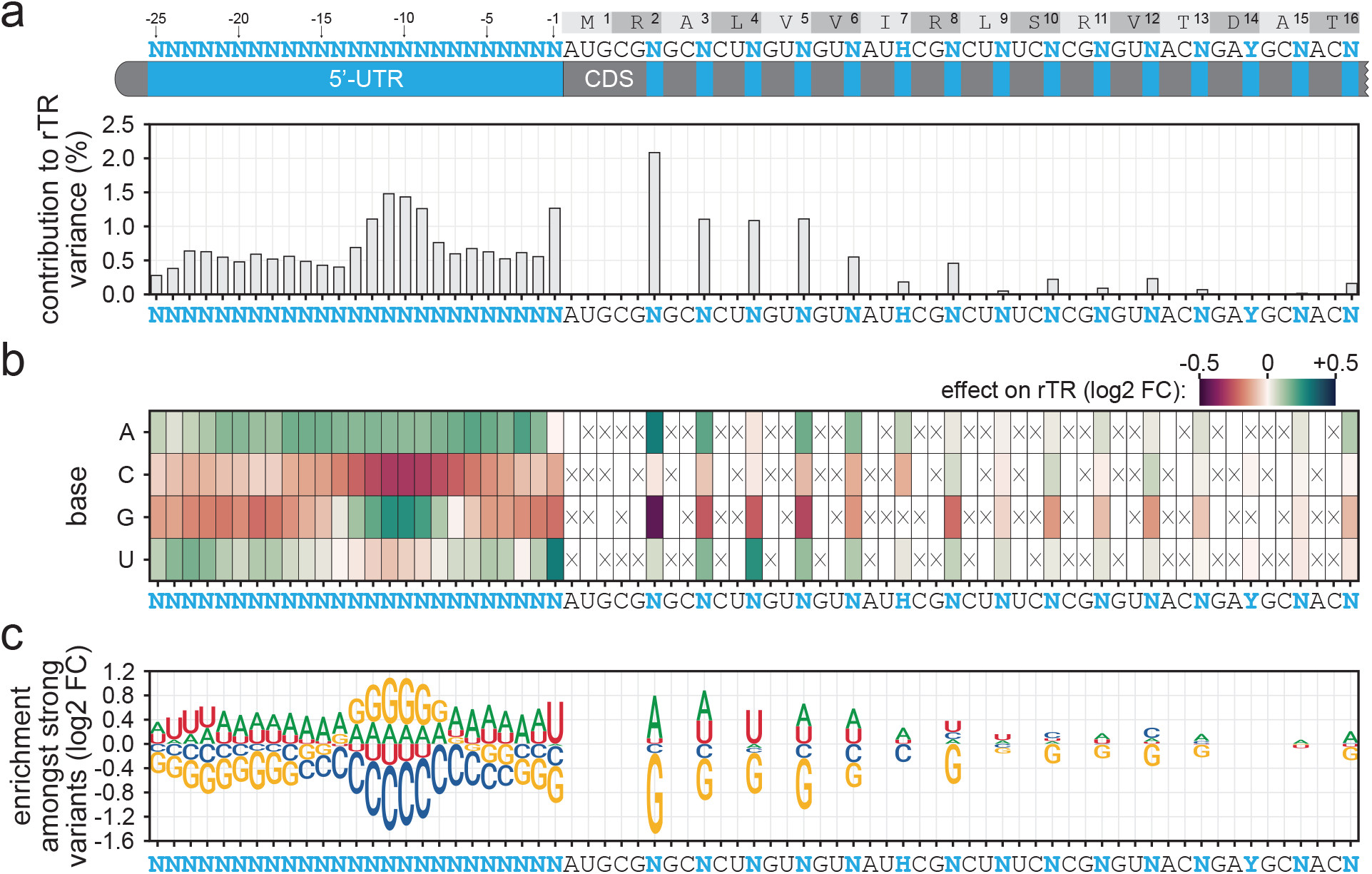
Positional and base-specific effects on translation. **a**) Contribution of variable mRNA positions to the observed rTR variance. The relative sum of squares calculated by ANOVA with each position as covariate is displayed. **b**) Base-specific effects of the randomised positions. Displayed effects are log2-transformed fold changes (log2 FC) of the mean rTR of variants with a given base at the respective position over the mean rTR of variants with any other base permitted at that position. Positive and negative values correspond to translation-increasing or -decreasing effects, respectively. Crossed boxes indicate non-permitted bases. **c**) Enrichment of bases amongst strong variants. The log2 FC of a base’s relative occurrence amongst strong variants (rTR ≥ 0.5) over its relative occurrence amongst weak variants (rTR < 0.5) is displayed.

### Quantification of sequence parameters and their effect on translation

Since less than 30% of variance in translation could be explained by global analysis of individual positions, we sought to examine the impact of different sequence parameters on the level of individual variants. Specifically, we computed several parameters known or hypothesised to influence rTR for all members of Lib_random_ and calculated their correlation with rTR. This analysis included parameters related to GC-content, hybridisation between mRNA and 16S rRNA, mRNA folding and other features. Since rTR values follow a non-normal distribution (p-value = 1.11 × 10^-79^, Shapiro-Wilk normality test) and some sequence parameters are likely to non-linearly correlate with rTR, we also report Spearman’s correlation (coefficient ρ) as a metric of rank correlation between parameters and rTR.

Overall GC-content shows significant correlation with the rTR (ρ^2^ = 18.6%, R^2^ = 11.3%) and its impact is higher in the 5’-UTR than the CDS (**Fig. 3a, Suppl. Fig. 11**). In particular high GC-content is strongly associated with low rTRs (**Suppl. Fig. 11**), likely due to a tendency of GC-rich sequences to form stable secondary structures, which are known to counteract translation (27). Further, we determined the minimum free energy (mfe), ensemble free energy (efe) and mRNA accessibility (acc) using two models for base pairing, the Turner energy model (T) and the CONTRAfold (C) model (65,66), resulting in six metrics related to mRNA folding: mfeT, mfeC, efeT, efeC, accT and accC (**Fig. 3a, Methods**). In brief, mfe and efe are energies required for the unfolding of the most likely and the ensemble of possible mRNA secondary structure(s), respectively, whereas acc is an accessibility score for a sliding window along the mRNA corresponding to the probability of this window being embedded within a secondary structure (18). Folding of mRNA showed a clear impact on rTR across all tested metrics (**Fig. 3a**). The latter show a positive correlation with the rTR, which is stronger than for GC-content and highest for efeC (ρ^2^ = 30.8%, R^2^ = 12.6%) and accC (ρ^2^ = 30.4%, R^2^ = 12.2%) (**Fig. 3a, b**). In particular very strong folding (e.g. efeC < −15 kcal × mol^-1^) completely abolishes efficient translation (**Fig. 3b**). We investigated further the impact of the positioning of secondary structures by calculating mRNA accessibility within a sliding window of ten nucleotides. Correlation of the resulting scores (accT/C_10nt_) with rTR is highest around the first few codons followed by the SD region, and sharply decreases further downstream in the CDS (**Fig. 3c**).

**Figure 3:**
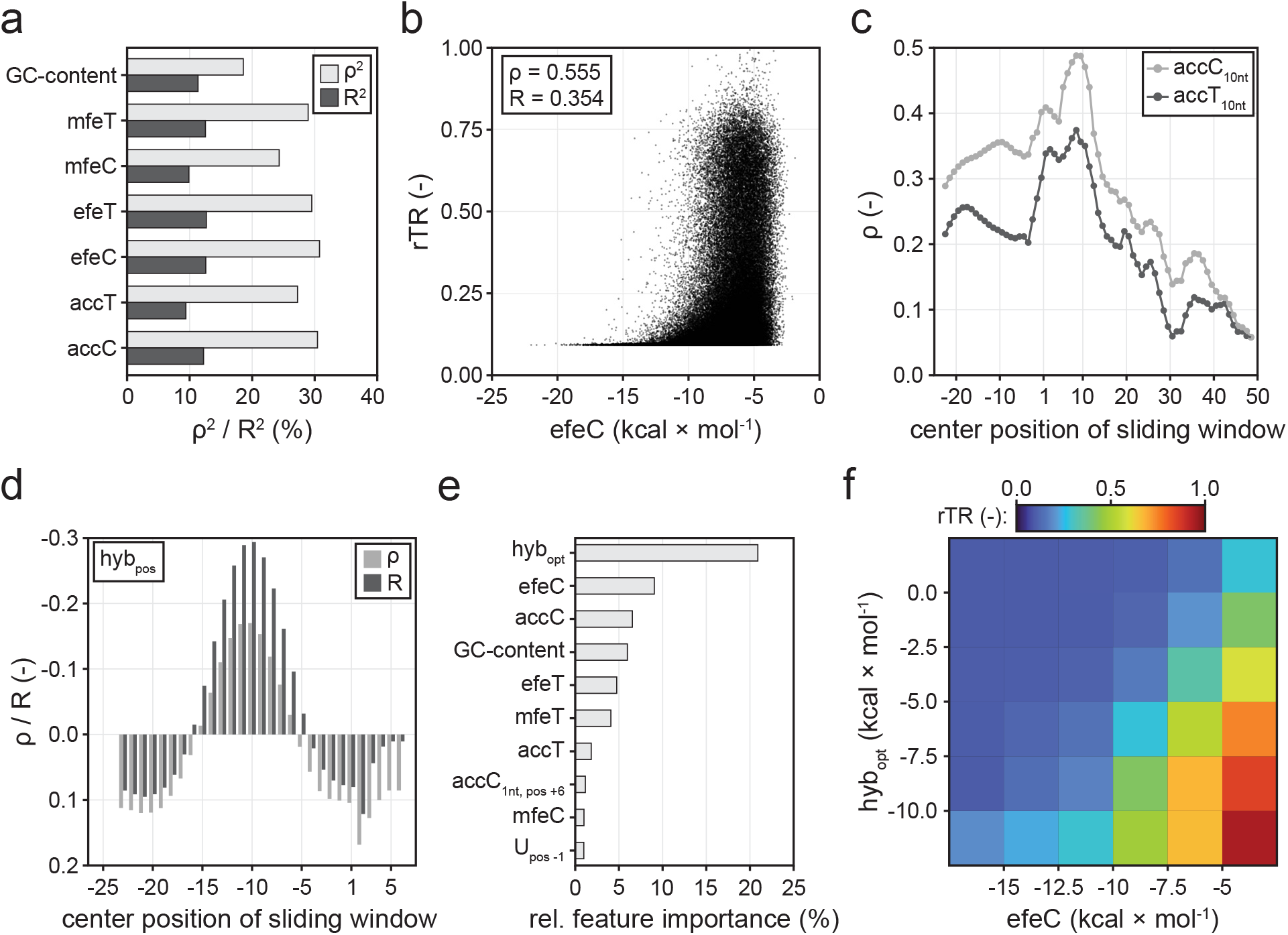
Effect of different sequence parameters on translation in Lib_random_. **a**) Correlation of GC-content and different mRNA folding metrics with rTR. Spearman’s ρ^2^ and Pearson’s R^2^ are displayed. **b**) Scatterplot between rTR and the best-correlating mRNA folding parameter efeC. **c**) Correlation of rTR with local mRNA accessibility. Parameters accT_10nt_ and accC_10nt_ correspond to the mRNA accessibility of a 10-nt window centered around the mRNA position specified on the horizontal axis. Endings C and T denote base pairing calculated by two different energy models (**Methods**). **d**) Correlation of hybridisation energy between 16S rRNA and different mRNA positions with rTR. Positional hybridisation energy (hyb_pos_) is displayed for 9-bp windows centered around the indicated mRNA position (horizontal axis). **e**) Relative feature importance of a random forest model trained on Lib_random_. The ten most important of 248 features are displayed. hyb_opt_: best-correlating hybridisation parameter (see main text). AccC_1nt, pos+6_: AccC score for position +6 of the mRNA. U_pos −1_: one-hot encoded U at position −1 of the mRNA. **f**) Mean rTR of variants in Lib_random_ as grouped by the two most predictive features of the random forest, hyb_opt_ and efeC. Tick labels mark the boundaries of the respective bins (boxes).

Next, we investigated the impact of interactions between mRNA and 16S rRNA. As expected, the hybridisation energy hybSD between *E. coli’s* 16S rRNA (sequence: 5’-ACCUCCUUA-3’) and the approximate SD region in the 5’-UTR (window between positions −18 and −4) shows a clear correlation with the rTR (**Suppl. Fig. 12**, **Methods**)(67). This observation is further corroborated by the fact that similarity with the canonical SD motif AGGAGGU in this window is strongly associated with high rTRs (**Suppl. Fig. 13**). Since the position of hybridisation is known to be critical for efficient translation, we further calculated positional hybridisation energies hyb_pos_ sliding the 9-nt 16S rRNA sequence along the mRNA (**Fig. 3d**, **Methods**). We found that hyb_pos_ is negatively correlated with rTR between 5’-UTR positions −15 and −6 indicating that stronger hybridisation (i.e. lower hyb_pos_) has a translation-favouring effect in this region. Outside of this window, a negative effect on rTR is observable. The 9-nt hybridisation window with the strongest correlation to rTR is centred around position −10 corresponding to a binding of the 16S rRNA 3’-end to the 5’-UTR between positions −14 and −6. A more systematic analysis of hybridisation windows and positions (**Suppl. Tab. 7**) revealed the mean of hybridisation energies at positions −11 and −10 (hyb_opt_) as the parameter with the highest correlation with rTR (ρ^2^ = 2.9%, R^2^ = 8.9%).

Based on those findings, we sought to quantify the utility of different sequence parameters for predictive modelling. To this end, we used the data from Lib_random_ to train a random forest regressor with the goal to predict the rTR from different features including primary sequence information as well as the above-mentioned secondary parameters (**Methods**). The model was trained using tenfold cross-validation (**Suppl. Fig. 14**) and its performance was evaluated on a test set strictly held out during training (randomly selected 10% of data). The resulting model predicts rTR values with good confidence (R^2^ = 58%, **Suppl. Fig. 15**). More importantly, we extracted the relative importance of features of the random forest (**Fig. 3e**). Remarkably, while the 16S rRNA hybridisation parameter hyb_opt_ had shown only moderate correlation coefficients ρ and R, it was by far the most important model feature (20.9%) followed by the folding parameters efeC (9.1%) and accC (6.5%). The over-proportional importance of hyb_opt_ could imply that successful hybridisation with the 16S rRNA must be fulfilled to obtain strong translation initiation rendering hyp_opt_ a critical, early decision criterion for the model. Furthermore, U at 5’-UTR position −1 ranked 10^th^ (1.0%) amongst the total of 248 encodings constituting the most important single-nucleotide feature. The majority of features (227) exhibited a relative importance below 0.5% pointing towards the multifactorial, interactive nature of the translation (initiation) process and likely to a high degree of redundancy between the tested encodings.

Lastly, we binned the variants from Lib_random_ according to the two most important features of the random forest, hyb_opt_ and efeC, and calculated the average rTR of each bin (**Fig. 3f**). Interestingly, we found that the appearance of very high rTRs (i.e. > 0.5) is co-dependent on strong 16S rRNA hybridisation and weak mRNA folding. Variants with strong secondary structures (efeC < −15 kcal × mol^-1^) only exhibit significant translation initiation if they hybridise well with the 16S rRNA. By contrast, variants with low folding energy can exhibit intermediate-to-strong translation even in the absence of SD motifs.

### Codon usage and interaction between 5’-UTR and CDS

A long-standing question is how strong the impact of the CDS on translation is, both in absolute terms and relative to the 5’-UTR. Changes in the CDS affect critical determinants of translation initiation such as codon usage and mRNA folding. Importantly, testing many different CDSs in combination with a single 5’-UTR (as amply done in previous studies) is insufficient to unambiguously assign observed effects to different sequence parameters and to quantify their contribution in a precise fashion, since some parameters also depend on and change with the 5’-UTR in place. Thus, it remains unclear if and how strong any observed effect is causally related to a sequence parameter change in a generalisable fashion, or whether it is merely a context-specific artefact only occurring for the selected 5’-UTR. Similarly, full randomization (as in Lib_random_ in this work) only delivers unique pairs of 5’-UTRs and CDSs, which again prohibits unambiguous attribution of effects to either of the two mRNA parts (5’-UTR or CDS). This problem can only be circumvented by testing large numbers of 5’-UTR-CDS combinations in a combinatorial manner with sufficient overlap allowing to average out case-specific artefacts. Therefore, to investigate the individual impact of 5’-UTR and CDS independently, we generated three additional libraries of combinatorial (Lib_comb1_, Lib_comb2_) and full-factorial (Lib_fact_) 5’-UTR-CDS pairs, which were constructed through combination of defined half-libraries (**Fig. 4a, Methods**): Lib_comb1_ combines about 1,000 5’-UTRs with about 1,000 CDSs, Lib_comb2_ is a combination of approximately 100 5’-UTRs with approximately 10,000 CDSs, and Lib_fact_ features ten independently cloned batches of about 100 5’-UTRs combined with about 100 CDSs each. Note that Lib_fact_ was designed such that in each batch every 5’-UTR is combined with every CDS and *vice versa* (i.e. full-factorial design). Next, we recorded the activity of variants from the three libraries applying the same uASPIre workflow as described for Lib_random_ above. Processing of NGS data yielded time series for 407,325, 496,643, 112,296 unique variants above high-quality read count threshold for Lib_comb1_, Lib_comb2_ and Lib_fact_, respectively. For Lib_fact_, two independent biological replicates were tested. We then grouped variants according to the 5’-UTR (or CDS) in place and analysed the diversity of the rTR amongst all CDSs (or 5’-UTRs) appearing with the respective fixed 5’-UTR (or CDS). Exchanging either 5’-UTR or CDS (while maintaining the other) can lead to strong up- and downshifts in expression (**Fig. 4b**). Shifts are on average much stronger for an exchange of the 5’-UTR than of the CDS, and in many cases cover a large fraction of the rTR range (**Fig. 4b, Suppl. Fig. 16**). We further quantified the individual impact of 5’-UTR and CDS performing an ANOVA with the mean rTRs of all 5’-UTRs and CDSs (**Fig. 4c, Methods**). This analysis was performed exclusively on Lib_fact_, since full-factorial design is required to exclude case-specific artefacts and achieve a precise quantification of each part’s individual contribution (see above). We observed that the 5’-UTR explains on average 53.12 ± 6.3% and the CDS 19.8 ± 5.4% of rTR variance confirming the higher impact of the 5’-UTR compared to the CDS. The 27.0 ± 1.3% of variance remain unexplained in the additive model and must therefore be caused by non-linear interactions between 5’-UTR and CDS confirming a high degree of interdependence between both parts.

**Figure 4:**
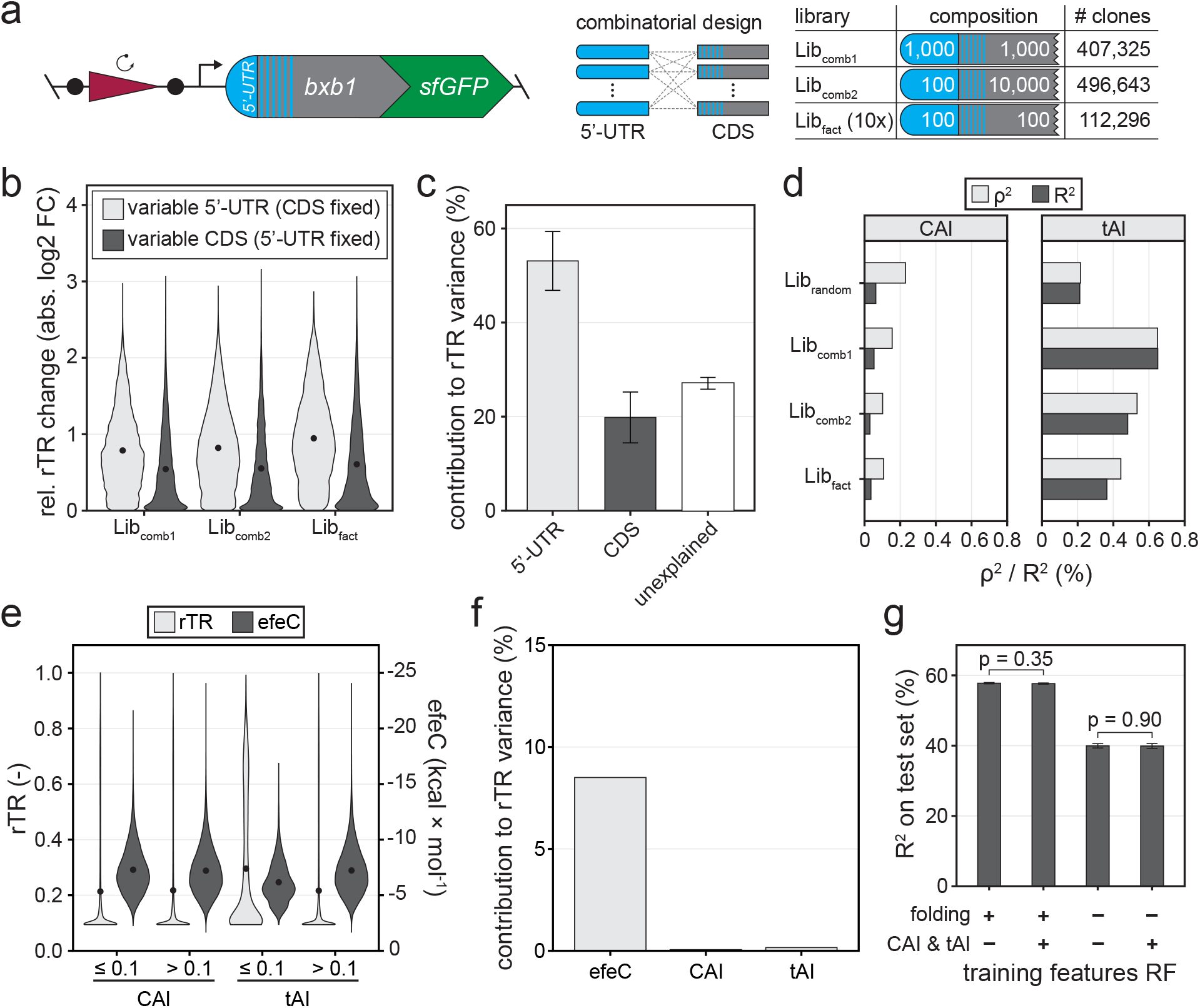
Overall impact of 5’-UTR, CDS and codon usage on translation. **a**) Three additional libraries of combinatorial (Lib_comb1_, Lib_comb2_) and full-factorial (Lib_fact_) design were assessed via uASPIre. Lib_comb1_: combinatorial combination of about 1,000 5’-UTRs and 1,000 CDSs. Lib_comb2_: combinatorial combination of about 100 5’-UTRs and 10,000 CDSs. Lib_fact_: ten independent batches, each a full factorial combination of approx. 100 5’-UTRs and 100 CDSs. Lib_fact_ was tested in two independent biological replicates. The number of analysed clones is indicated for each library. **b**) Impact of the exchange of 5’-UTRs or CDSs on translation. The rTR change (absolute value) of a given 5’-UTR upon exchanging its CDS versus the mean rTR of all variants with that same 5’-UTR is displayed (and *vice versa*). Black circles within violins are mean relative rTR changes. **c**) ANOVA with the mean rTRs of all 5’-UTRs and CDSs in Lib_fact_. Error bars: standard deviation between ten independent batches of Lib_fact_. **d**) Correlation of codon usage indices CAI and tAI with rTR. **e**) Comparison of rTRs and folding energies (efeC) of variants with low (≤ 10%) and high (> 10%) CAI/tAI in all libraries. Black circles within violins are mean rTR/efeC values. **f**) Contribution of efeC, CAI and tAI to the rTR variance in all libraries according to an ANOVA with only the three parameters as covariates. **g**) Impact of folding and codon usage metrics on the performance of random forest (RF) models trained on Lib_random_. Sequence parameters for mRNA folding (mfeT, mfeC, efeT, efeC, accT and acc) and codon usage (CAI and tAI) were added or omitted during training. Error bars: Standard deviation of five training repeats with 10fold cross-validation each. p-value were calculated with Welch two sample t-tests.

A controversially discussed sequence feature of the CDS is codon usage, which is well known to influence translation (initiation). To this end, the appearance of rare codons within the first few triplets of the CDS was found to coincide with high expression (29,41–44). Thus, we first analysed the impact of two commonly used metrics for codon usage, CAI and tAI (**Suppl. Tab. 6**) (41,53), on rTR, which indicated a weak (R^2^ and ρ^2^ consistently below 0.7%) yet significant correlation in all libraries (**Fig. 4d**). However, it remains unclear whether this is caused by differential abundance of the corresponding tRNAs in the cell or by changes in mRNA folding. Since folding is also co-dependant on the 5’-UTR in place, combinatorial testing of 5’-UTR-CDS pairs is also essential in this case to unambiguously test if and to which extent the two aforementioned hypotheses are correct. Accordingly, we first compared the rTR of Lib_fact_ variants rich in rare codons (i.e. CAI/tAI ≤ 10%) with the other variants (i.e. CAI/tAI > 10%). Variants with low CAI exhibit a mean rTR of 0.213, which is virtually indifferent from high-CAI variants (0.217) (**Fig. 4e**). This is further corroborated by the fact that the mean rTRs of CDSs and CAI do not correlate significantly (p-value = 0.256, one sample t-test) in Lib_fact_ (**Suppl. Fig. 17**). Low-tAI variants, by contrast, exhibits on average a higher rTR than the control group (**Fig. 4e, Suppl. Fig. 17**). At the same time, however, mRNA folding is significantly weaker (p-value < 10^-300^, one-sided Welch two sample t-test) in low-versus high-tAI variants, which is not the case for the corresponding CAI groups (p-value = 1.0, **Fig. 4e**). Moreover, the codon frequency of *E. coli* showed only very small and inconsistent effects on the rTR for the randomised codons (**Suppl. Fig. 18**). Therefore, we further analysed to which extent the dependence of the rTR on codon usage can be explained by mRNA folding. An ANOVA with only efeC, CAI and tAI as covariates indicated that the overwhelming majority of variance in rTR explainable by these parameters is attributed to efeC (8.5%), whereas the contribution CAI and tAI was about 155- and 53-fold lower, respectively (**Fig. 4f**). Furthermore, we re-trained the former random forest model (see above) with different sets of sequence parameters including CAI and tAI (**Fig. 4g**). Remarkably, while removal of mRNA folding parameters led to a substantial decrease in model performance, addition of CAI and tAI did neither increase accuracy of the initial random forest nor was it able to compensate for the performance loss in the absence of folding parameters. Accordingly, the relative feature importance of CAI and tAI was very low (**Suppl. Fig. 19**). Collectively, these findings strongly suggest that any influence of codon usage on rTR can be virtually completely explained by mRNA folding. On the contrary, a causal connection to cellular tRNA abundance or the previously postulated translational ramps could not be established and is either insignificant or negligible amongst the over 1.2 million sequences tested in this study.

### Assessment of translational anomalies of arginine codon 2 and 5’-UTR position −1

Lastly, we sought to decipher the reasons for the unexpected behaviour of two mRNA positions observable in our data (see above). To this end, the presence of G in the third position of arginine codon 2 and U in position −1 of the 5’-UTR exhibit a profound impact on the rTR, which is negative in the former and positive in the latter case (compare **Fig. 2**). Variants with CG<u>G</u> as the second codon show an average decrease in rTR of 26.3% compared to variants carrying A, C or U in the third position (**Fig. 5a**). This different behaviour is likely not caused by codon frequencies or tRNA availability, since both arginine codons with higher (CG<u>C</u>, CG<u>U</u>) and lower (CG<u>A</u>) frequency show significantly higher mean rTRs (**Fig. 5b**). By contrast, the average folding energy of CG<u>G</u>-bearing variants is significantly lower (△efeC = −0.76 kcal × mol^-1^) than for the other codons (**Fig. 5a**), pointing again to mRNA folding (and not tRNA availability) as the mechanistic reason for the differential expression of synonymous codons. For variants with U at position −1 in the 5’-UTR, the mean rTR is 23.3% higher than for those with any other base in this position (**Fig. 5c**). In this case, however, the average folding energy is even slightly increased for U (△efeC = +0.10 kcal × mol^-1^) excluding mRNA folding as the reason (**Fig. 5c**). As an alternative explanation, we suspected that an interaction of this U with the initiator tRNA (tRNA^fMet^) could be responsible for the observed effect. In *E. coli*, initiator tRNAs are encoded by one monocistronic (*metY*) and one tricistronic (*metZWV*) transcriptional unit, and their sequences are identical except for position 46 (G in *metY*, A in *metZWV*). Importantly, methionine elongator tRNAs (*metT, metU*) do not initiate translation (71), and all tRNA^fMet^ copies carry an A in position 37 directly 3’ to the CAU anticodon, which could preferentially hybridise with mRNAs carrying a U directly 5’ to the start codon.

**Figure 5:**
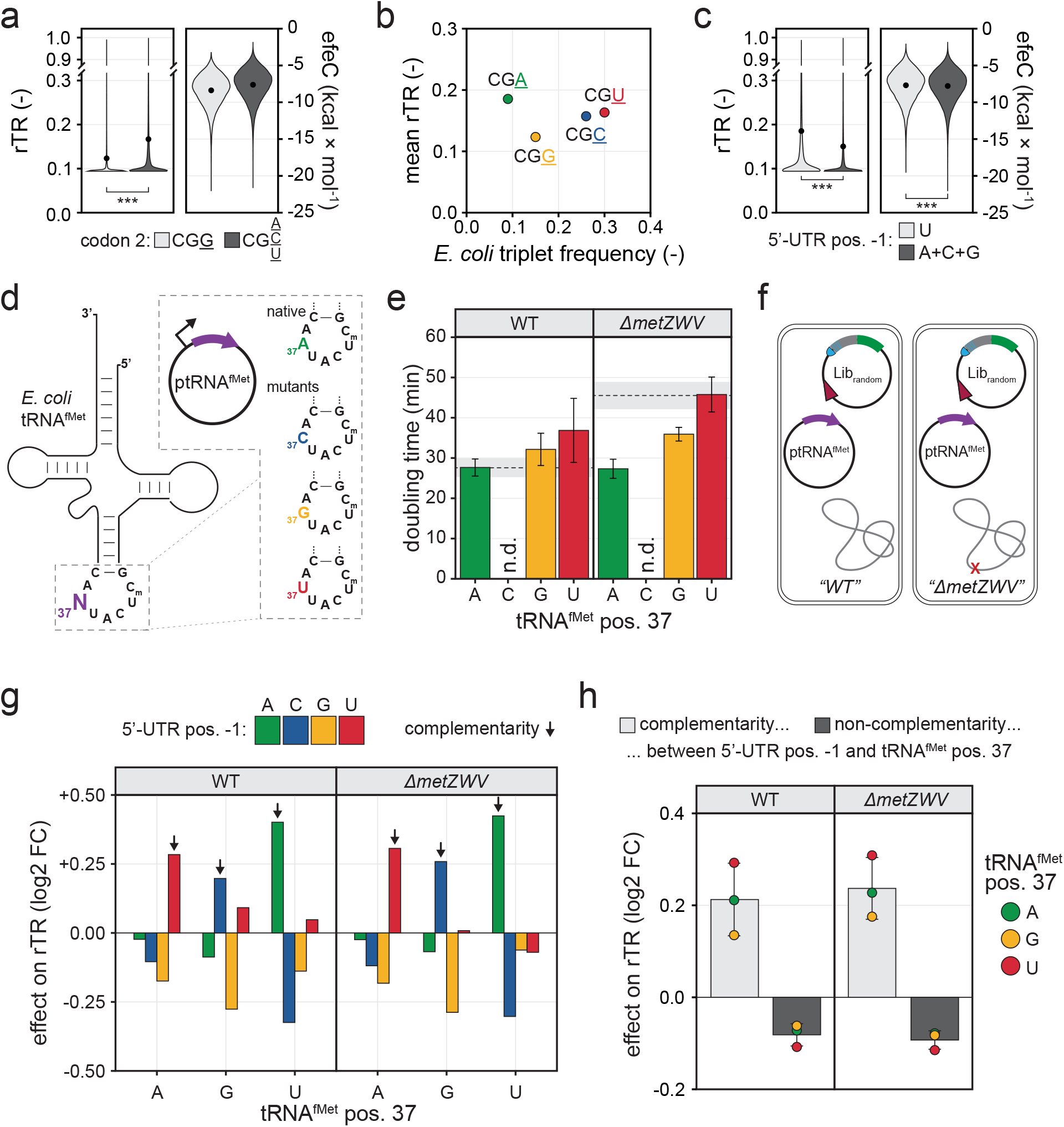
Assessment of translational anomalies of arginine codon 2 and 5’-UTR position −1 in Lib_random_. **a**) Effect of different synonymous codons in the second triplet of the CDS on rTR and mRNA folding energy (efeC). Black circles within violins are mean rTR/efeC values. *** denote p-values <10^-16^ in a Welch two sample t-test. **b**) Relationship between relative triplet frequency in *E. coli* and rTR for the four synonymous triplets in arginine codon 2. **c**) Effect of different bases in 5’-UTR position −1 on rTR and mRNA folding energy (efeC). Black circles within violins are mean rTR/efeC values. *** denote p-values<10^-16^ in a Welch two sample t-test. **d**) Plasmids for the overexpression of native initiator tRNA^fMet^ and mutants thereof (**Suppl. Fig. 6, Methods**). Position 37 (3’-adjacent to the CAU anticodon) of tRNA^fMet^ is mutated from A to C, G, or T/U. **e**) Growth of *E. coli* strains carrying plasmids for tRNA^fMet^ overexpression in shake flask cultivations (LB, 37 °C). Bars are mean doubling times of independent biological triplicate cultivations with standard deviation as error bars. Dashed lines are the mean doubling time of the respective strain without tRNA overexpression (i.e. empty vector control) with standard deviation as grey shaded areas. For tRNA^fMet-A37C^, doubling times were not determined (n.d.) due to severe growth inhibition (see main text). **f**) Approximately 50,000 variants of Lib_random_ were tested in the presence of overexpressed tRNA^fMet^ variants in *E. coli* strains containing (WT) and lacking (*ΔmetZWV*) the chromosomal *metZWV* locus. **g**) Impact of tRNA^fMet^ mutations on the rTR of variants from Lib_random_. Displayed effects are log2-transformed fold-changes (log2 FC) of the average rTR of variants with a given base at 5’-UTR position −1 over the average rTR of variants with any other base at this position. Black arrows indicate complementarity between 5’-UTR position −1 and position 37 of the tRNA^fMet^ variant. **h**) Impact of complementarity between 5’-UTR position −1 and tRNA^fMet^ position 37. Circles are log2-transformed fold-changes (log2 FC) of the average rTR of variants with complementarity or non-complementarity between mRNA and tRNA over the mean rTR of all variants in the same group (i.e. same tRNA^fMet^ variant and strain). Bars are the mean log2 FCs of the three tRNA^fMet^ variants for each case and strain with standard deviation as error bars.

Several previous studies have postulated or shown that the presence of U in this position favours formation of the prokaryotic ribosomal initiation complex and/or translation of the corresponding genes *in vitro* and *in vivo* (36,58,72-80). These effects were attributed to a proposed interaction of A37 in tRNA^fMet^ and U in 5’-UTR position −1, for which further evidence was later provided in algal chloroplasts through compensatory mutation of tRNA^fMet^ position 37 (81). Furthermore, structural analyses have shown that A37 is released from internal base pairing upon reaching the ribosomal P-site (82), which would render this position available for Watson-Crick base pairing with nucleotide(s) upstream of the start codon. Collectively, these prior works highlight the importance of bases directly upstream of the start codon and point to a potential interaction of mRNA and tRNA^fMet^ beyond the codon-anticodon hybridisation. A causal link between any observed impact on translation and an interaction with the 5’-UTR position −1 was, however, so far not conclusively established. A potential reason for this could be that only few mRNA sequence variants were tested prohibiting generalisable statements due to the high context dependence of translation initiation and statistical error.

We therefore investigated whether the proposed interaction between mRNA and tRNA^fMet^ could be substantiated relying on systematic high-throughput sequence-function mapping. We first constructed plasmids for the overexpression of tRNA^fMet^ with the native A37 as well as the mutants A37C, A37G and A37U (**Fig. 5d, Methods**). To reduce the background from the chromosomal tRNA^fMet^ copies, we further deleted the *metZWV* locus of *E. coli* TOP10 *ΔrhaA* (“WT”) yielding strain TOP10 Δ*rhaA ΔmetZWV*(“*ΔmetZWV*”), and transformed both strains with the tRNA plasmids. Note that simultaneous knockout of *metZWV* and *metY* failed in our hands despite complementation via plasmid-borne tRNA^fMet^. Remarkably, transformants of ptRNA^fMet-A37C^ showed severe growth inhibition (colonies visible only few days after transformation), whereas the native tRNA^fMet-A37^ and the other two mutants (tRNA^fMet-A37G^, tRNA^fMet-A37U^) were tolerated with minor effects on growth in both strains (**Fig. 5e**). While in the case of the WT strain a small increase of doubling times was observable, *ΔmetZWV* showed an improvement of growth upon overexpression of all tRNA^fMet^ variants, likely due to compensation of the reduced level of chromosomally-derived tRNA^fMet^ copies in this strain. The apparent toxicity of tRNA^fMet-A37C^ could stem from global dysregulation of translational, and due to its prohibitively slow growth we excluded this variant from further experiments. Next, we tested approximately 50,000 variants from Lib_random_ in both strains (WT, *ΔmetZWV*) in presence of the remaining tRNA^fMet^ plasmids via uASPIre (**Fig. 5f**, **Suppl. Fig. 20**, **Methods**). We analysed the resulting NGS data comparing 44,289 common 5’-UTR-CDS variants above high-quality read count threshold that appeared in all six conditions (i.e. two strains with three plasmids). Specifically, we determined for each condition the effects of 5’-UTR position −1 by dividing the mean rTR of variants with a given base at this position by the mean rTR of all other variants (**Fig. 5g**). This analysis confirmed the strong, base-specific impact of this position, and, beyond that, revealed a significant dependence of the effect on the base present in position 37 of tRNA^fMet^. To this end, we observed a strong increase in the rTR for variants whose base upstream of the start codon is complementary to position 37 of the overexpressed tRNA^fMet^ in both the WT and *ΔmetZWV* strain. Non-complementarity consistently leads to a lower expression compared to the complementarity case across both strains and all tRNA^fMet^ variants (**Fig. 5h**). Similarly, a small yet significant positive impact on rTR is observable for the major wobble base pair G-U/U-G, which appears consistently for both directions of interaction (G in position 37 of tRNA^fMet^ with U in 5’-UTR position −1 and *vice versa*) and both strains (**Suppl Fig. 21**). Interestingly, a U at 5’-UTR position −1 leads to a small rTR-boosting effect also in presence of the non-complementary initiators tRNA^fMet-A37G^ and tRNA^fMet-A37U^ only in the WT strain (**Fig. 5g**). This can be explained by the presence of chromosomally encoded, endogenous tRNA^fMet-A37^ copies, since this positive effect is neutralised or slightly inverted in the *ΔmetZWV* strain. The effects at all other randomised positions in the mRNA were similar to the ones obtained for Lib_random_ without overexpression of tRNA^fMet^ variants (**Suppl. Fig. 22** compare **Fig. 2b**).

These findings strongly suggest a direct base-pairing interaction of 5’-UTR position −1 with the nucleotide following the anticodon in tRNA^fMet^ (position 37), which leads to a significant positive effect on translation initiation upon successful hybridisation. Thus, our analyses confirm previous hypotheses to that end in a statistically solid manner based on more than 132,000 mRNA-tRNA^fMet^ combinations, which were kinetically assessed in two different genetic backgrounds.

## DISCUSSION

In this study, we systematically investigated the impact of 5’-UTR and N-terminal CDS on translation through mapping of more than 1.2 million mRNA sequence variants to their corresponding expression levels in *E. coli*. In combination with random and combinatorial library design, the ultrahigh throughput of our approach allowed us to critically assess sequence parameters known or supposed to influence translation efficiency. Furthermore, the generated large data basis enabled a precise quantification of effect sizes and correction for sequence-specific artefacts via statistically solid analyses.

To this end, we assessed mean effects of individual bases and positions in 5’-UTR and CDS along with various higher-order sequence parameters of the mRNA. We found that 25% of variance in our data could be explained by individual nucleotides and that GC-content, hybridisation with the 16S-rRNA and mRNA folding are the most significant determinants of translation confirming findings from previous studies (e.g. (8–12,20,27–30,32–37,39,40,83)). Using a simplistic machine learning approach, we compared the predictive potential of 248 parameters, which ranked 16S-rRNA hybridisation highest (20.9%) followed by various mRNA folding features (between 1.0% and 9.1%) and GC-content (5.97%) and pointed to a high degree of interaction and redundancy amongst parameters (**Fig. 5e**).

Furthermore, we found an unexpectedly large, base-specific contribution of two individual nucleotides, the negative impact of G in the third position of arginine codon 2 and the positive effect of U in position −1 of the 5’-UTR (**Fig. 2**). Follow-up analyses revealed that the former is not causally related to tRNA availability in the cell but can likely be attributed to a stronger tendency of variants with CG<u>G</u> as second codon to form mRNA secondary structures (**Fig. a, b**). Notably, mRNA accessibility at this position ranked amongst the most important features (accC_1nt_, _pos. +6_) of a predictive random forest model (**Fig. 3e**), which confirms the relation of the observed effect to mRNA folding. The positive effect of U directly upstream of the start codon, by contrast, was not linked to folding or any other mRNA parameter (**Fig. 5c**), which prompted further experiments to that end. Specifically, we assessed whether a base-pairing interaction of 5’-UTR position −1 with the base in 3’ to the anticodon in initiator tRNA^fMet^ (position A37) could be responsible for the effect. This hypothesis could be confirmed through compensatory mutation of tRNA^fMet^ position 37, which led to a translation-favouring effect in all cases of complementarity between tRNA and 5’-UTR (**Fig. 5g, h**). Several previous studies had shown that a U upstream of the start codon favours ribosome assembly and/or translation *in vitro* and *in vivo* (36,58,72–80). A link of these effects to the aforementioned base-pairing interaction, however, was only postulated and not experimentally confirmed in these studies. Esposito *et al*. (81) attempted to confirm the interaction in algal chloroplasts by substitution of position A37 in tRNA^fMet^. Notably, the variant tRNA^fMet-A37C^ was not generated in their study, which showed severe growth inhibition and was thus excluded also in our work. For the tested reporter gene (*petA*), substitution of A37 indeed led to a translation-favouring effect only in cases of complementarity between tRNA^fMet^ and the base upstream of the start codon. However, this observation was made on the basis of only three 5’-UTR position −1 variants of *petA* carrying a non-native weak UAA start codon and could not be confirmed for several other analysed genes. Whether the impact for *petA* is specific to this gene (context) or the weak start codon, or indeed related to an interaction between tRNA^fMet^ and the base upstream of the start codon therefore remains unclear. In this study, we assessed 45,258 mRNA sequences tested with three tRNA^fMet^ variants and in two strains of normal and reduced endogenous expression of native tRNA^fMet-A37^. This did not only confirm the proposed quadruplet interaction in a statistically firm fashion but allowed to even quantify comparably subtle phenomena such as wobble base pairing (**Suppl. Fig. 21**), which can be masked for individual sequences and thus are inaccessible to low-throughput approaches.

Lastly, we constructed and assessed more than a million combinatorial and full-factorial 5’-UTR-CDS combinations, which, in view of the high degree of interactivity, is indispensable to correctly assign observed effects to different mRNA parts and sequence parameters, and to precisely measure their contribution. This allowed us to quantify the mean individual contribution of the 5’-UTR and CDS to translational variance in a manner that would not be possible otherwise (e.g. using fully random libraries), which amount to 53% and 20%, respectively. Moreover, we capitalised on the combinatorial libraries and the large data basis to revise different hypotheses on the causal relationship between translation efficiency and codon usage. Similar to previous studies (e.g. (33,37)), our data confirmed a strong dependence of the rTR on the N-terminal CDS and a decreasing impact of codons with increasing distance to the start codon (**Figs. 2, 4b, c**). While this dependence unquestionably exists, the underlying mechanistic reasons remain less clear and were linked to both differences in mRNA folding and cellular tRNA abundance in the past. In our data, we found a small (R^2^/ρ^2^ < 0.7%) yet significant correlation of the rTR with codon usage metrics (**Fig. 4d**). However, the majority of the corresponding variance of the rTR can be explained by mRNA folding to an overwhelming degree while the contribution of codon usage metrics is extremely low (**Fig. 4f**). This low impact is further corroborated by the fact that none of the codon usage metrics was capable to increase the prediction accuracy of a random forest model, whereas mRNA folding had a very large impact (**Fig. 4g**). In summary, amongst the 1.2 million unique 5’-UTR-CDS combinations tested in this study the influence of different codons is virtually fully explainable by mRNA folding, whereas a causal connection to cellular tRNA abundance was either insignificant or negligibly small. The small apparent correlation between codon usage indices and rTR thus likely stems from differences in GC-content between rare and frequent codons, which leads to different tendencies to form secondary mRNA structures.

Consequently, our study highlights the importance of ultradeep sequence-function mapping for the accurate determination of the contribution of parts and phenomena involved in gene regulation. It should be mentioned that several other factors are known to influence translation (initiation), which have not been addressed in this study. These include the use of different start codons, long-range interactions between ribosome and 5’-UTR, and limitations of translation elongation (e.g. related to protein folding). Nonetheless, the presented methodology can be applied to scrutinise these additional factors, which, together with the results from this study, could serve as a basis to improve on inaccuracies of currently available models for the prediction and forward design of prokaryotic protein expression.

## ACKNOWLEDGEMENT

We thank Dr. Christian Beisel (ETH Zurich) for his support with NGS, and Dr. Nico Claassens and Dr. Thijs Nieuwkoop for critical reading of the manuscript.

## AUTHOR CONTRIBUTIONS

M.J. and S.H. conceived the study and planned experiments. S.H. performed experiments and computational works. S.H. and M.J. analysed data. M.J. coordinated the study. S.H. and M.J. wrote the manuscript.

## FUNDING

This work was supported by the European Commission [grant number 766975], and the Swiss National Science Foundation under the NCCR “Molecular Systems Engineering”.

## CONFLICT OF INTEREST

The authors declare no conflict of interest.

## Notes

### Competing Interest Statement

The authors have declared no competing interest.

