## Supplementary Information for "Ultradeep characterisation of translational sequence determinants refutes rare-codon hypothesis and unveils quadruplet base pairing of initiator tRNA and transcript"

### 1. Supplementary figures

All DNA sequences follow the IUPAC nucleotide code; N: A/C/G/T; H: A/C/T; Y: C/T.

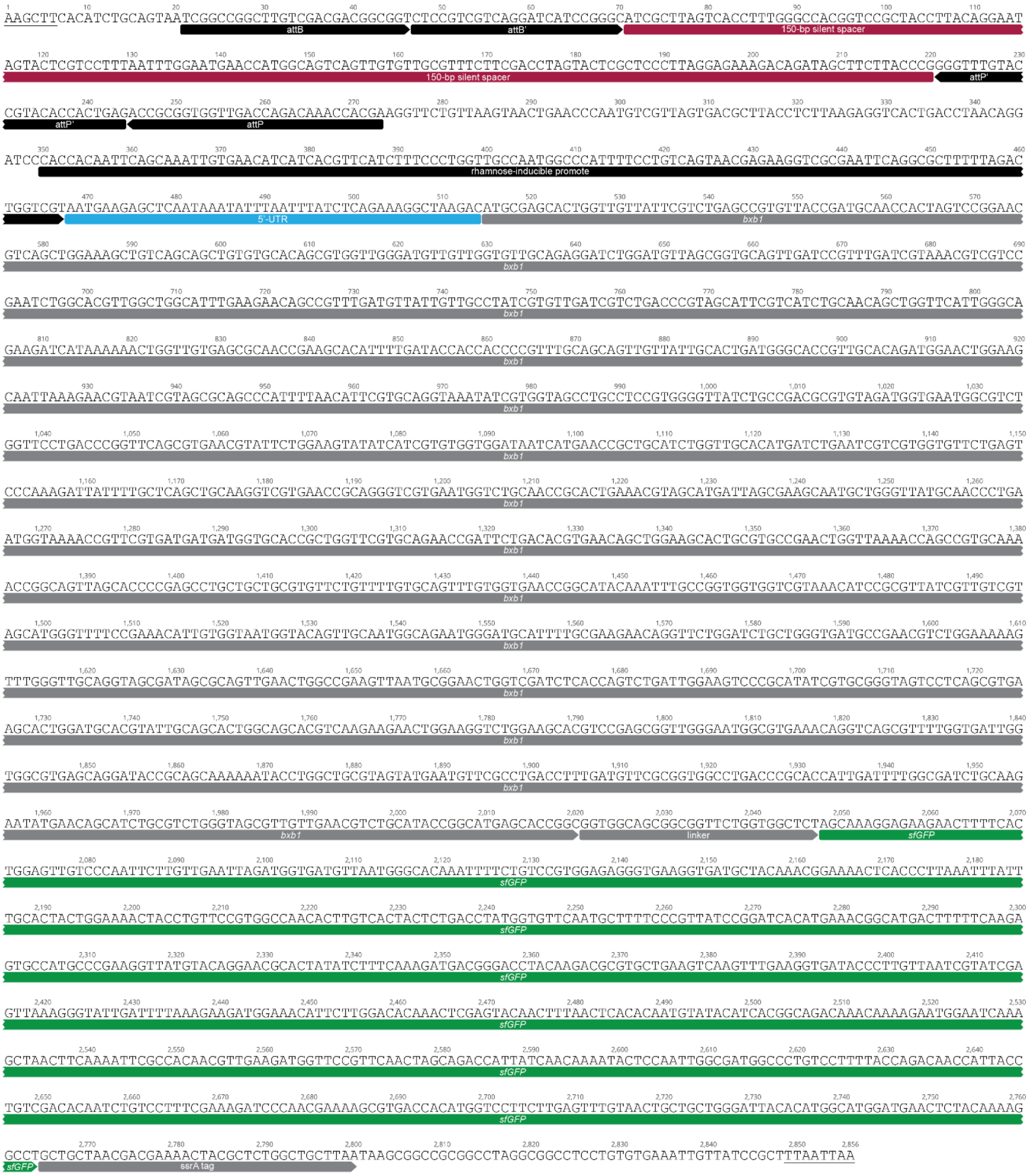


**Supplementary Figure 1: Graphical representation of pASPIre4**. The displayed sequence corresponds to the insert between HindIII and PacI restriction sites (underlined) in pSEVA291. ssrA: proteolytic degradation tag. The full sequence is available in text format in **Supplementary Note 2**.

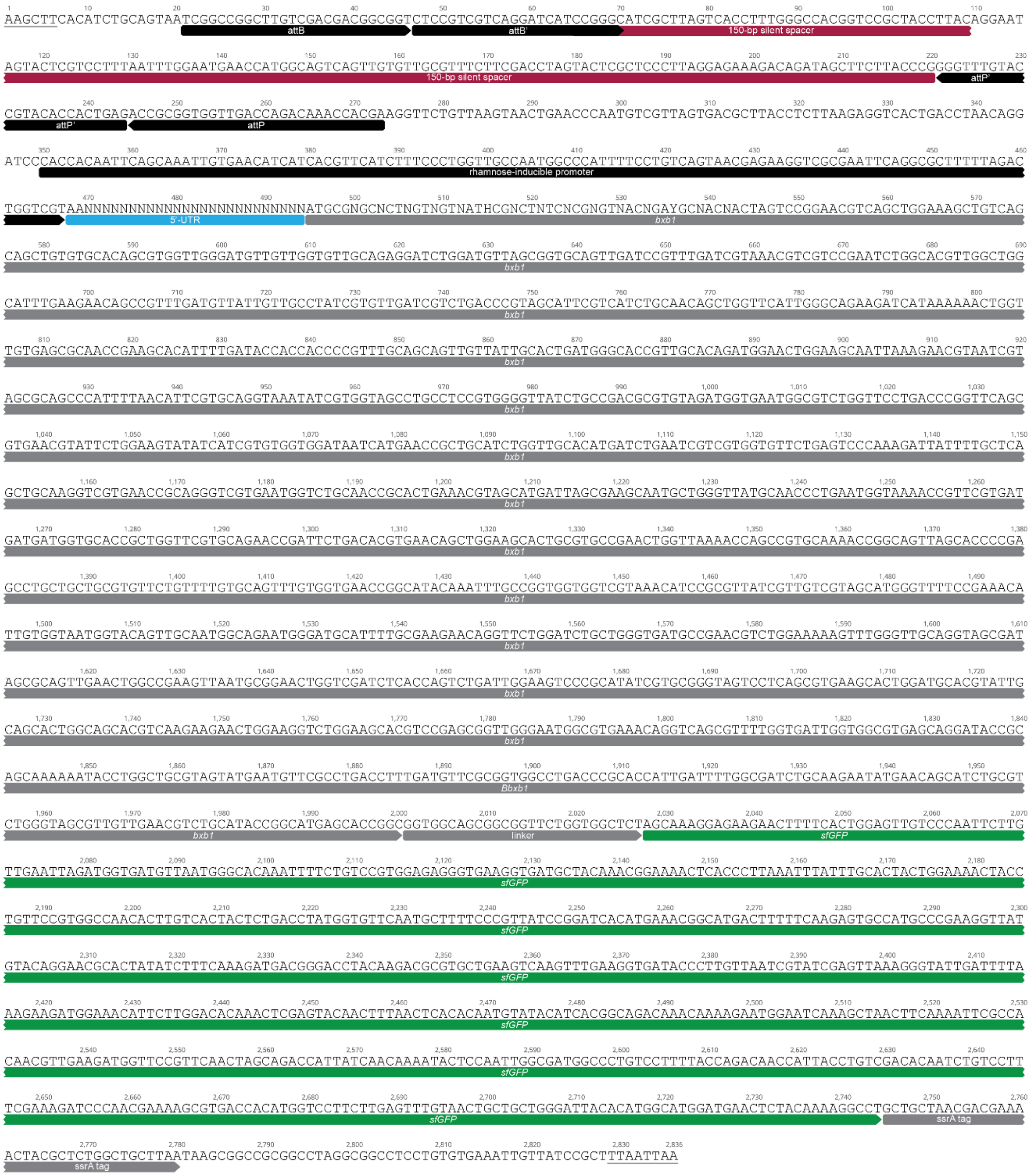


**Supplementary Figure 2: Graphical representation of pASPIre4$_{\mathbf{lib}}$.** The displayed sequence corresponds to the insert between HindIII and PacI restriction sites (underlined) in pASPIre4. ssrA: proteolytic degradation tag. The full sequence is available in text format in **Supplementary Note 3.**

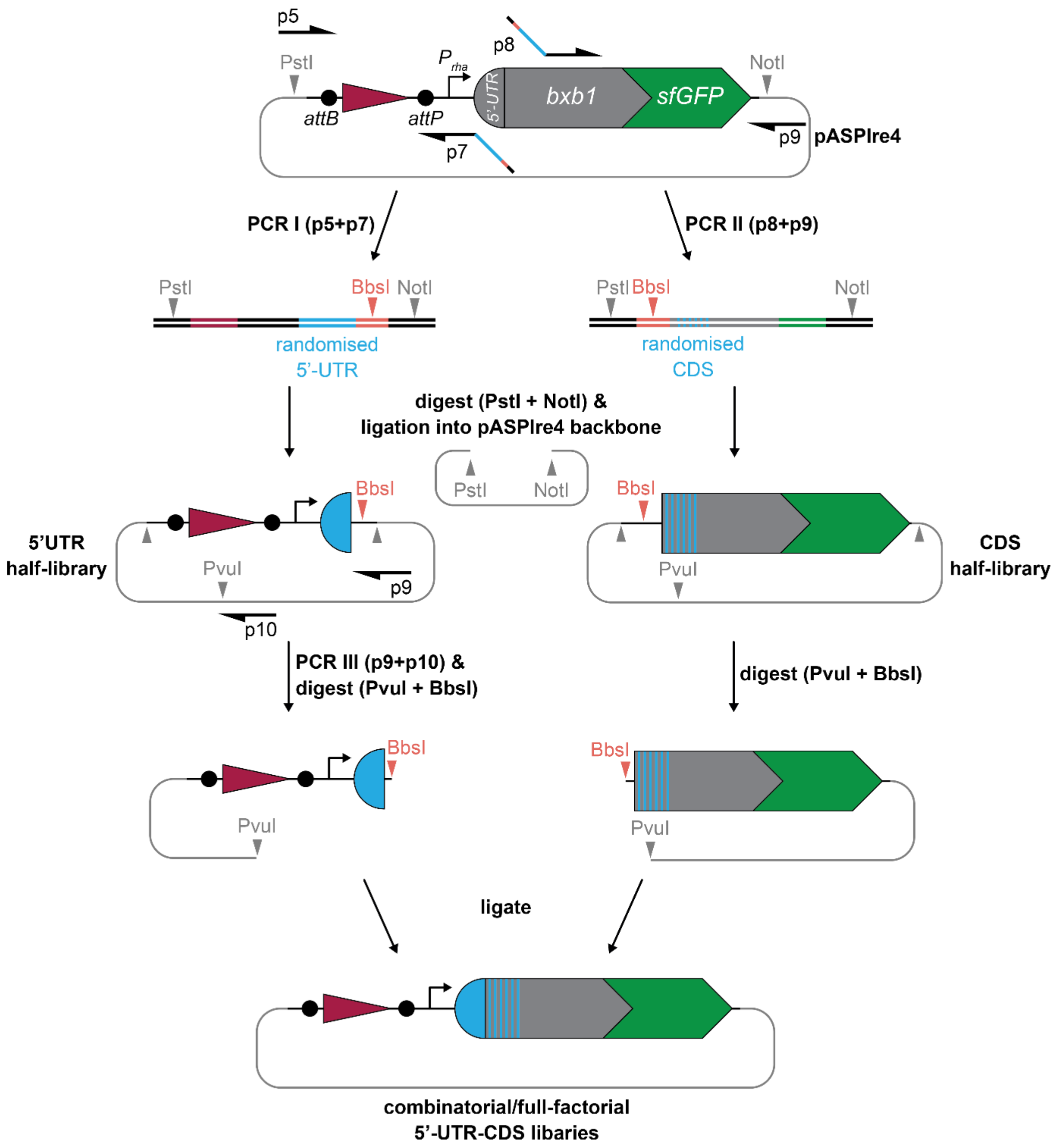


**Supplementary Figure 3: Cloning scheme for the generation of combinatorial and full-factorial 5'-UTR-CDS libraries.** Libraries were generated from 5'-UTR and CDS half-libraries as described in the **Methods** section. The 5'-UTR (left) half-library was generated by PCR with primers p5 and p7 with pASPIre4 as template. Primer p7 introduces degeneracy in the 5'-UTR and a BbsI site between the randomised 5'-UTR and the NotI site. The CDS half-library (right) was generated by PCR with primers p8 and p9 on pASPIre4 as template. Primer p8 introduces degeneracy in the CDS and a BbsI site between the CDS and the PstI site. The resulting PCR products were then sub-cloned into pASPIre4 to retrieve half libraries. In a second step, 5'-UTR and CDS half-libraries were combined to generate full libraries $Lib_{comb1}$, $Lib_{comb2}$ and $Lib_{fact}$. To achieve this, plasmid DNA of the 5'-UTR half-library was PCR-amplified with primers p9 and p10, and the product was digested with BbsI and PvuI and ligated into the backbone prepared from the CDS half-library via digestion with PvuI and BbsI. Note that the BbsI type IIS restriction site enables scarless joining of 5'-UTR and CDS half-libraries using ATGC (start codon ATG + first downstream base) as sticky ends for ligation.

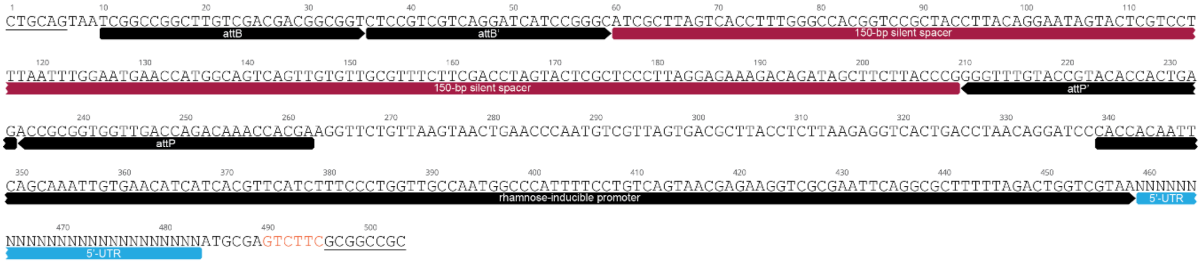


**Supplementary Figure 4: Graphical representation of 5’-UTR half-libraries**. The displayed sequence corresponds to the insert between PstI and NotI restriction sites (underlined) in pASPIre4. The introduced BbsI restriction site (orange) is used for scarless joining of 5’-UTR and CDS half-libraries. The full sequence is available in text format in **Supplementary Note 4**.

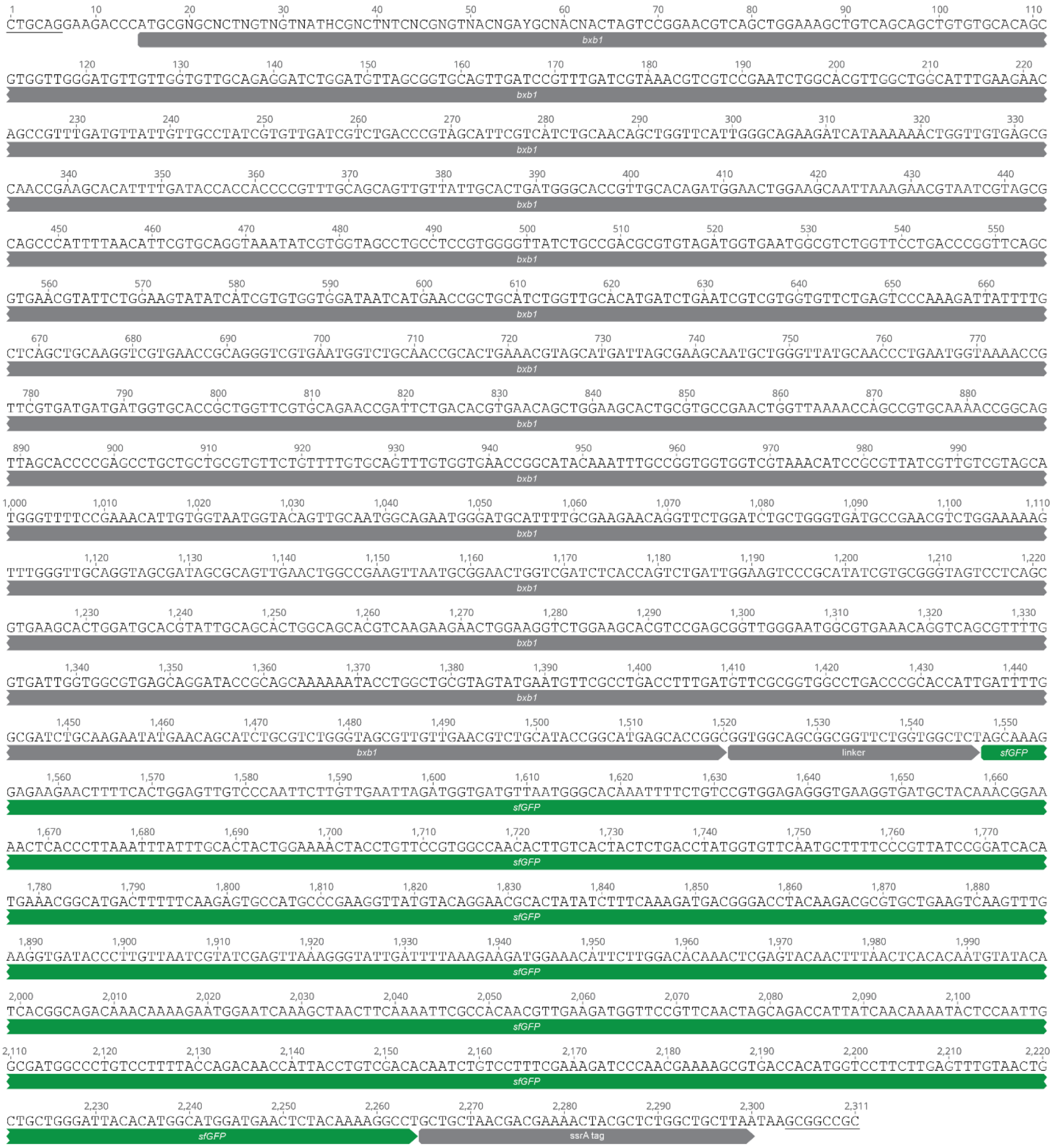


**Supplementary Figure 5: Graphical representation of CDS half-libraries.** The displayed sequence corresponds to the insert between PstI and NotI restriction sites (underlined) in pASPIre4. The introduced BbsI restriction site (orange) is used for scarless joining of 5’-UTR and CDS half-libraries. The full sequence in text format is available in **Supplementary Note 5.**

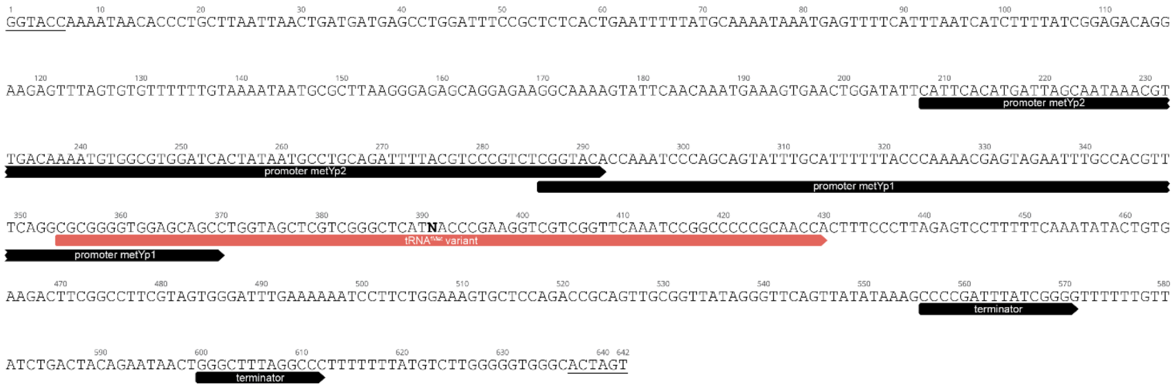


**Supplementary Figure 6: Graphical representation of plasmids for the overexpression of tRNA$^{fMet}$ variants**. The displayed sequence corresponds to the native chromosomal locus of *metY* including regulatory sequences. The bold N nucleotide indicates the mutated position 37 (p37) in tRNA$^{fMet}$. The window between KpnI and SpeI restriction sites (underlined) in pSEVA361 is shown. The full sequence is available in text format in **Supplementary Note 6**.

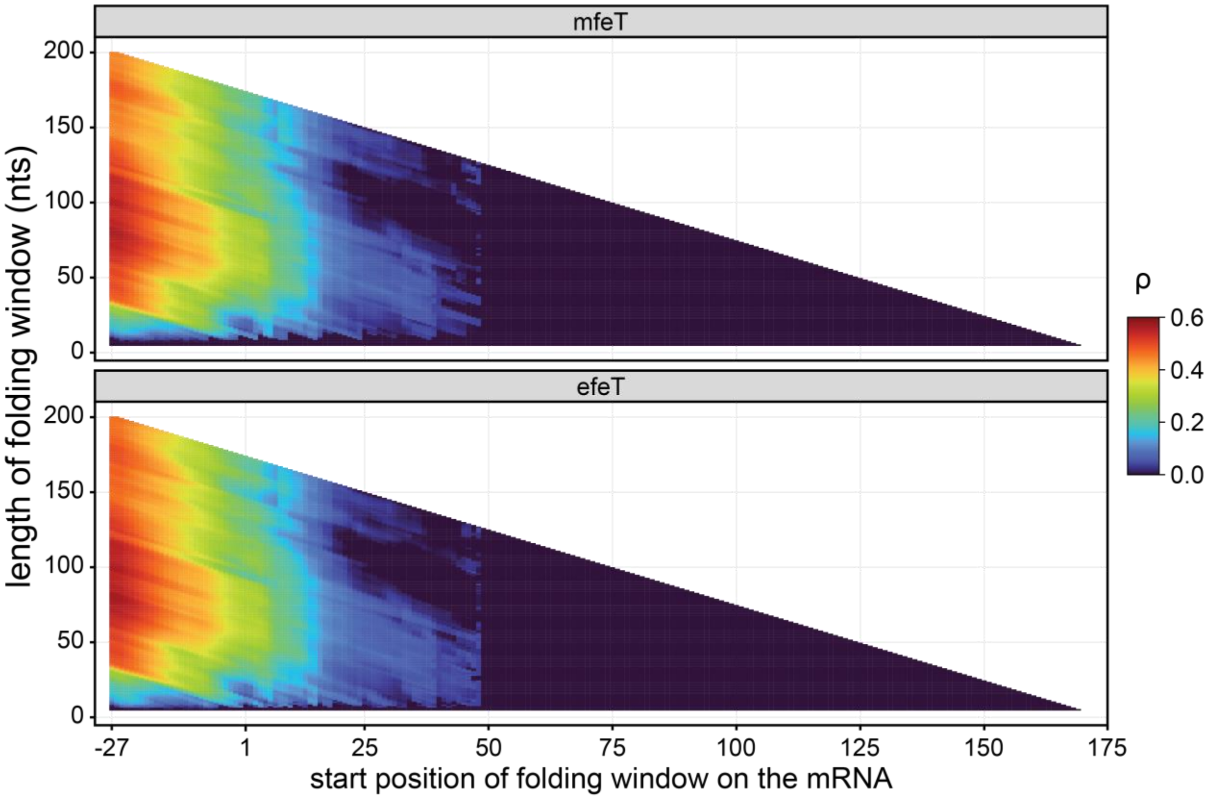


**Supplementary Figure 7: Correlation between folding energy and rTR for different mRNA windows**. Folding energies (mfeT and efeT) of all possible mRNA sequence windows with lengths between 5 and 200 bases were calculated using *RNAfold* of the *Vienna RNA* package and the resulting values were correlated with rTR using Spearman's correlation (**Methods**).

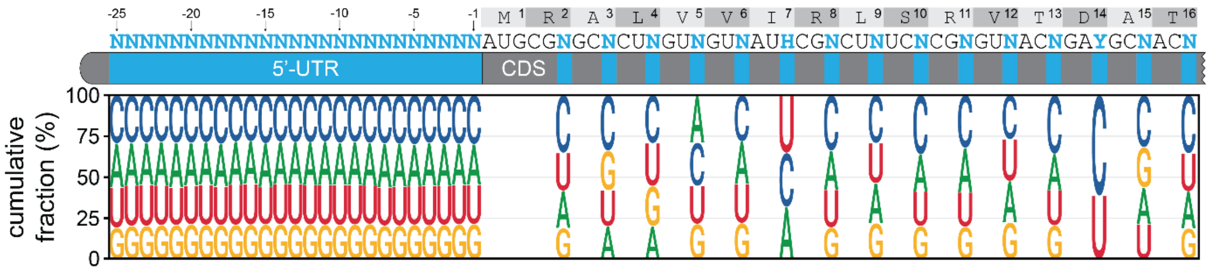


**Supplementary Figure 8: Base distribution in Lib$_{random}$.** The base distribution amongst all 198,174 variants of Lib$_{random}$ above high quality read count threshold is displayed for each randomised position. Bases are ordered from the most (top) to the least (bottom) frequent for each position. Note that in codons 7 (Ile) and 14 (Asp) only three (A, C, U) and two (C, U) bases were allowed, respectively, to avoid non-synonymous mutations.

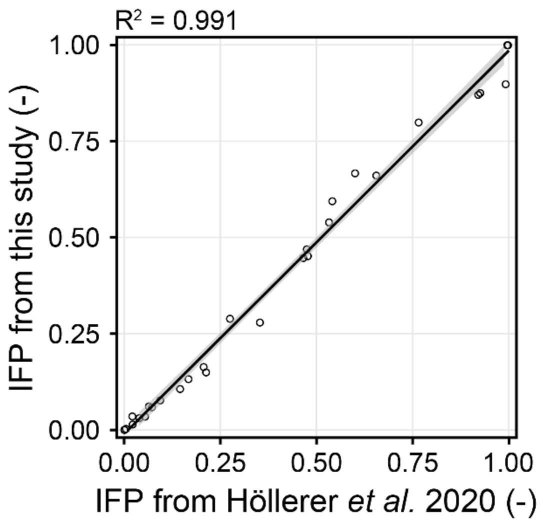


**Supplementary Figure 9: Cross-study reproducibility of uASPIre.** The IFP of 31 standard RBSs as measured in this study is plotted over the IFP of the same standard RBSs as determined in a previous study (1). The black line corresponds to a linear regression with 95% confidence interval (grey area).

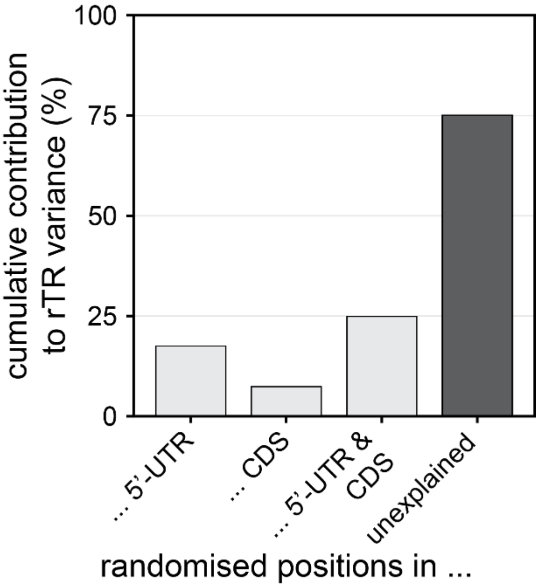


**Supplementary Figure 10: Cumulative impact of randomised positions in 5'-UTR and CDS on the rTR.** Bars are the cumulative contribution of all randomised positions in the mRNA part(s) indicated on the horizontal axis except for "unexplained" which corresponds to the residuals. The contribution of individual positions (**Fig. 2a**) was determined by ANOVA (**Methods**).

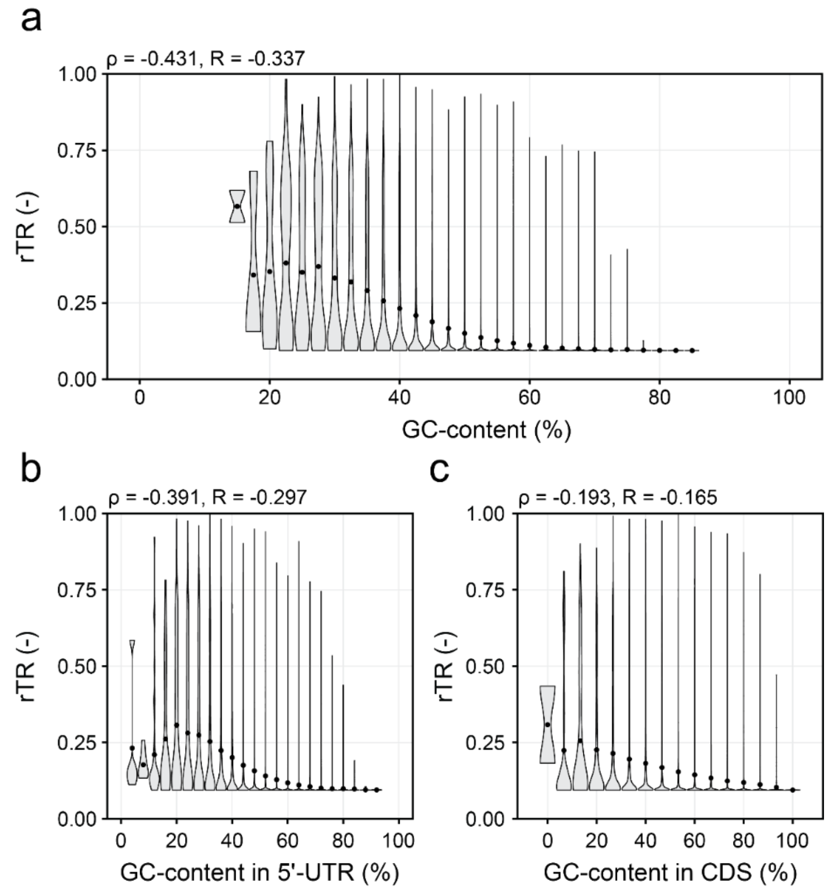


**Supplementary Figure 11: Impact of GC-content on rTR.** The correlation of rTR and GC-content in all randomised positions (**a**) as well as in the randomised positions in 5'-UTR (**b**) and CDS (**c**) individually was assessed. Variants were binned according to their GC-content in the respective mRNA part, and the rTR distribution of bins is displayed as violin with each bin's mean rTR (black circles).

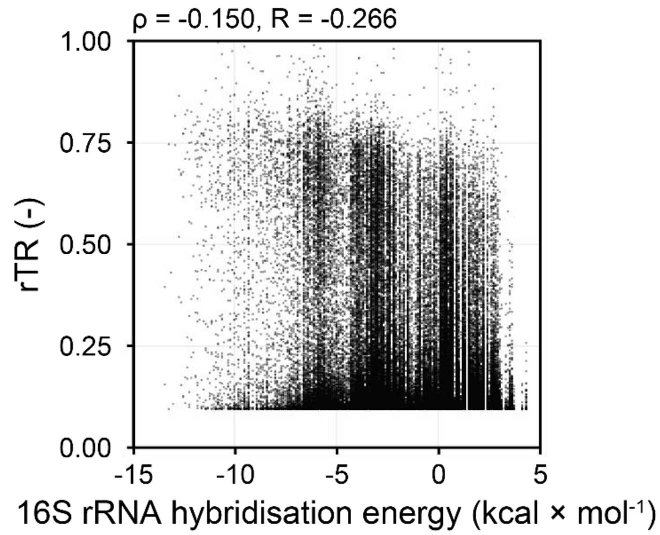


**Supplementary Figure 12: Correlation between 16S rRNA hybridisation energy and rTR**. The hybridisation energy of the 3'-end of *E. coli*'s 16S rRNA (sequence: 5'-ACCUCCUUA-3') and the 5'-UTR between position -18 and -4 was calculated using *RNAduplex* of the *Vienna RNA* package and the resulting energy values were correlated with rTR.

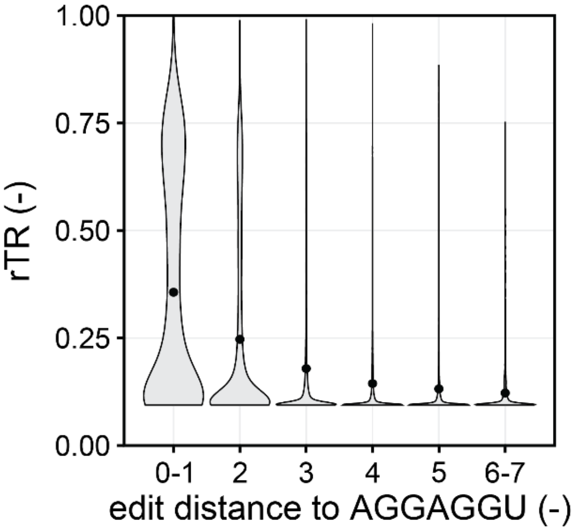


**Supplementary Figure 13: Impact of similarity to the SD motif on rTR**. The minimum edit distance between the canonical SD motif AGGAGGU and all 7-nt windows between 5'-UTR positions -18 and -4 was calculated for each variant (**Methods**). Violins are rTR distributions amongst variants grouped by minimum edit distance with group means (black circles).

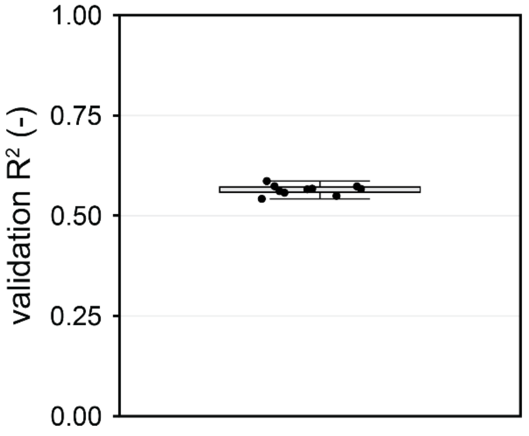


**Supplementary Figure 14: Cross-validation of random forest model (Fig. 3e)**. Validation $R^2$ values are shown for 10 independent validation runs as circles. Box represents interquartile range with whiskers marking the 1.5-fold interquartile range.

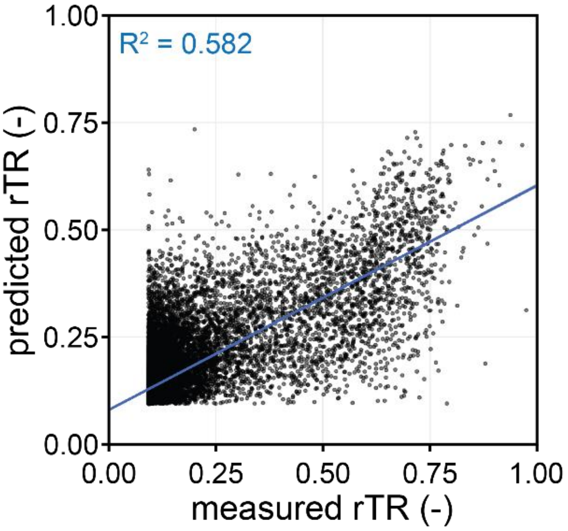


**Supplementary Figure 15: Random forest performance.** The rTR values predicted by the random forest model trained on $Lib_{random}$ (compare **Fig. 3e**) are plotted over the experimentally measured rTRs for 19,816 randomly selected test set variants strictly held out during training. The blue line corresponds to a linear regression with corresponding Pearson's $R^2$.

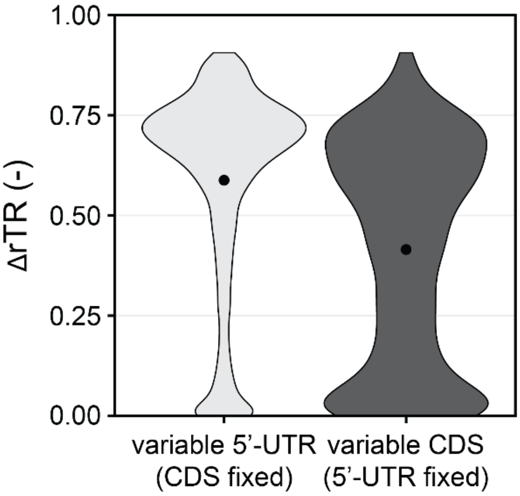


**Supplementary Figure 16: Coverage of rTR range upon exchange of 5’-UTR (left) and CDS (right).** For each CDS (or 5’-UTR) that appeared at least twice in all libraries, the absolute difference in rTR between the strongest and weakest 5’-UTR (or CDS) combined with this CDS (or 5’-UTR) is displayed (ΔrTR). Violins indicate the distribution of ΔrTR with black circles representing the mean value.

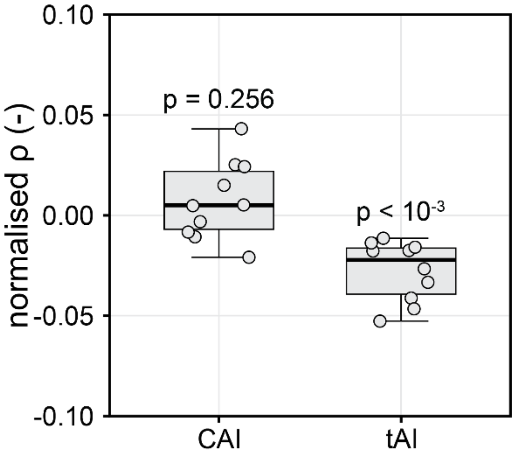


**Supplementary Figure 17: Correlation between codon indices and the CDS-derived contribution to rTR variance**. Pearson’s correlation between CAI/tAI and the mean rTR of each CDS (over all 5’-UTRs combined with that CDS) was calculated for each of the ten batches of $Lib_{fact}$ (grey circles) and normalised to the mean effect of the CDS on rTR (Fig. 4c). Boxes represents interquartile range with whiskers marking the 1.5-fold interquartile range and medians as solid black lines. P-values of one sample t-tests are indicated above each box.

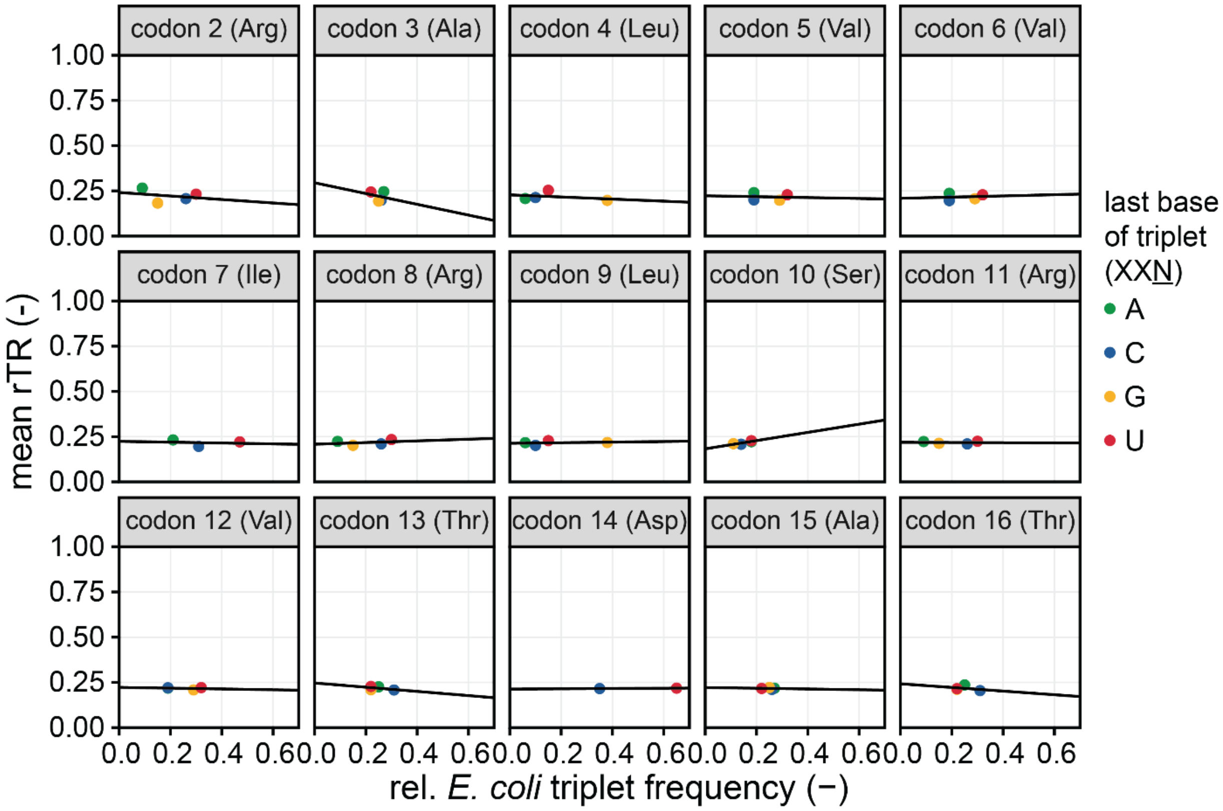


**Supplementary Figure 18: Dependence between triplet frequency in *E. coli* and rTR.** Mean rTRs of variants from all libraries with the same triplet in the randomised codons are plotted over the relative codon frequency of the respective triplet in *E. coli*. Triplet frequencies can be found in **Supplementary Table 6**. Black lines are best linear fits of data points. Note that no consistent up- or downward trend is observed across all codons as well as amongst triplets coding for the same amino acid but in different positions.

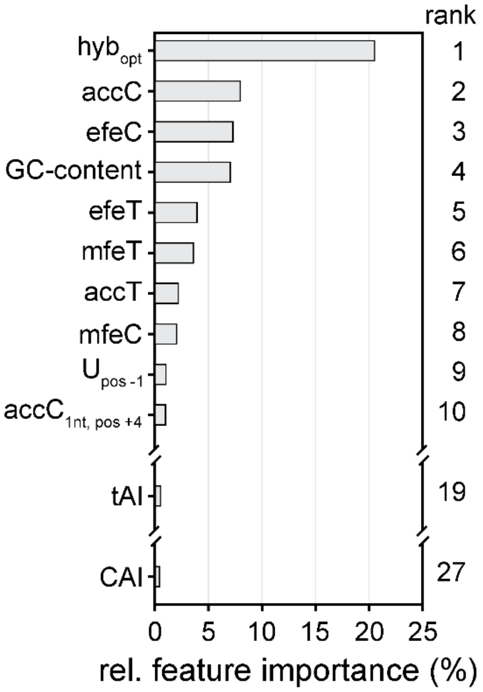


**Supplementary Figure 19: Feature importance of random forest model trained with codon usage and folding metrics.** Relative importance of the ten most important features as well as tAI and CAI for the model displayed in **Figure 5g** (second bar from left) is shown. Ranking of features is indicated to the right. $AccC_{1nt,\ pos\ +4}$: AccC score for position +4 of the CDS.

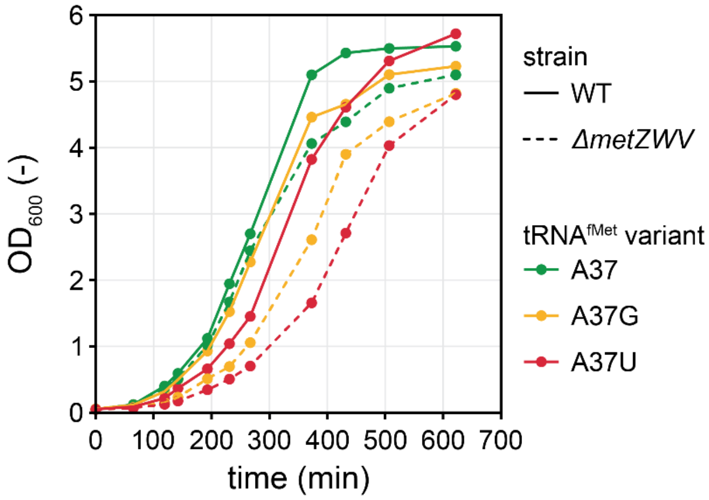


**Supplementary Figure 20: Growth of *E. coli* WT and *ΔmetZWV* expressing different $tRNA^{fMet}$ variants.** Approximately 50,000 variants of $Lib_{random}$ were co-cultivated in shake flasks (**Methods**) for each of the six combinations of *E. coli* strains (WT or *ΔmetZWV*) and overexpressed $tRNA^{fMet}$ variants (A37, A37G or A37U).

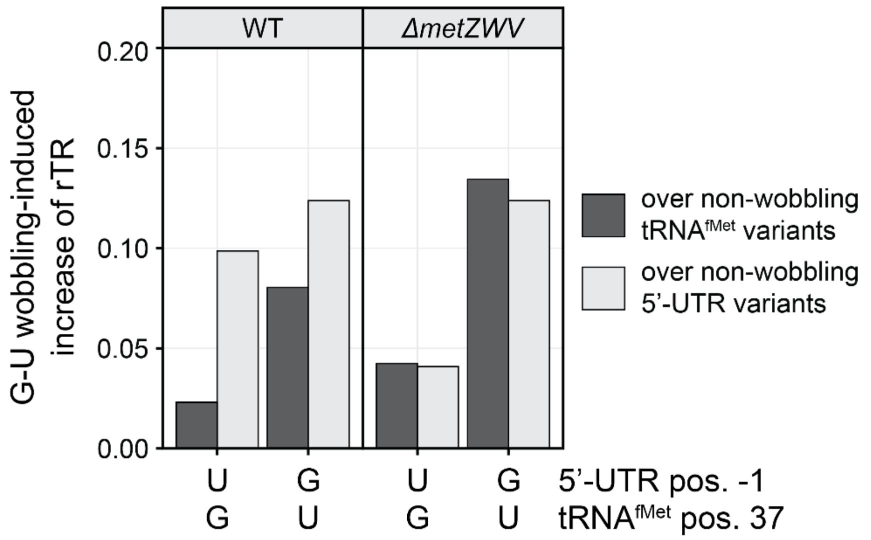


**Supplementary Figure 21: Effect of G-U wobbling on rTR.** The mean increase of the rTR is shown for different combinations of G-U wobbling base pairs between 5'-UTR position -1 and $tRNA^{fMet}$ position 37 as indicated on the horizontal axis. The rTR increases of the respective wobbling pair over either all non-wobbling and non-complementary $tRNA^{fMet}$ variants (dark grey) or over all non-wobbling and non-complementary 5'-UTR variants (light grey) are displayed. In order to facilitate comparison between the different experimental groups (i.e. six combinations of strain and $tRNA^{fMet}$ variant), rTR values were normalised to the mean rTR of each group prior to determination of the G-U wobbling-induced increase of rTR.

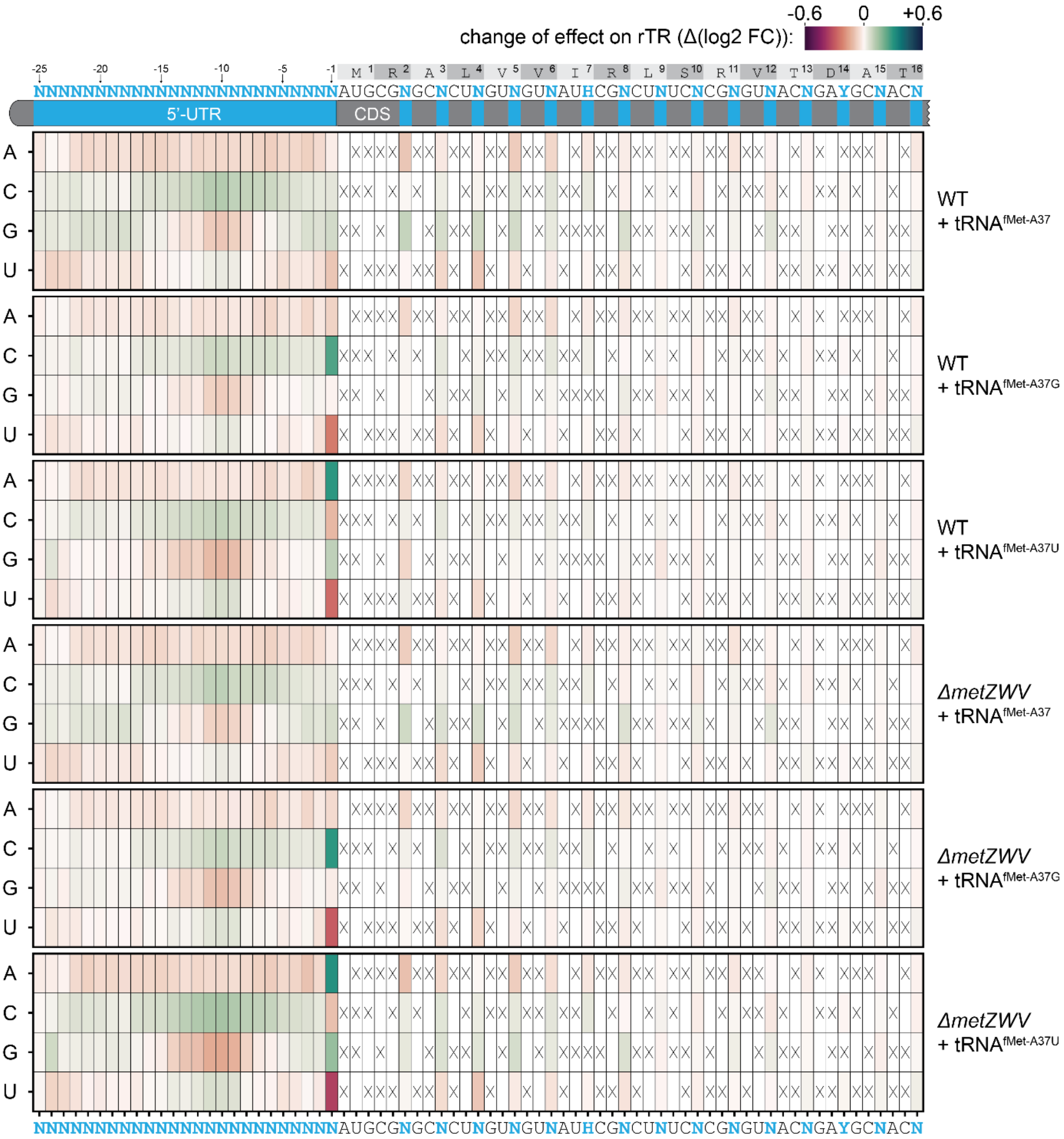


**Supplementary Figure 22: Base- and position-specific change of the rTR upon overexpression of tRNA$^{fMet}$ variants.** The change of the effect on rTR compared to the case without tRNA overexpression in the WT strain (**Fig. 2b**) is shown for different combinations of strains (WT vs. Δ*metZWV*) and overexpressed tRNA$^{fMet}$ variants (A37, A37G or A37U). This change of effect was calculated as follows: First, log2-transformed fold changes (log2 FC) of the mean rTR of all variants with a given base at the respective position over the mean rTR of all variants with any other base permitted at that position were determined for variants of Lib$_{random}$. Afterwards, the log2 FCs obtained for the WT strain without tRNA$^{fMet}$ overexpression (**Fig. 2b**) were subtracted (i.e. Δ(log2 FC)). Crossed boxes indicate non-permitted bases.

### Supplementary tables

**Supplementary Table 1: Primers used in this study.** Restriction sites (BbsI/NotI/PstI/SpeI) are underlined. The introduced BbsI site is marked in orange. Primer binding regions are highlighted in bold.

| Name | Sequence (5'-3') | Description |
|---|---|---|
| p1 | TTTGTTCAAAATCATGCCAAATCCGTGATCGGGGTAAAAAA**TGATCGGCACGTAAGAGGTTCC** | Forward primer for the generation of the knockout cassette *ΔmetZWV::specR* |
| p2 | GAGAAGGGGATGATAAAAAGGCGCTGAATGGCGCTTTTTTA**TTATTGGCTGGCACCAAGCAG** | Forward primer for the generation of the knockout cassette *ΔmetZWV::specR* |
| p3 | **GCGGCAAGCATTGCCACAACCGTGC** | Forward genotyping primer for *ΔmetZWV* |
| p4 | **CGTATTTTGCCGATGGGGCGACGCTGG** | Reverse genotyping primer for *ΔmetZWV* |
| p5 | **TCTATCAACAGGAGTCCAAG** | Forward primer for the generation of all libraries |
| p6 | TATATA<u>ACTAGT</u>NGTNGCRTCNGTNACNCGNGANAGNCGDATNACNACNAGNGCNCGCATNNNNNNNNNNNNNNNNNNNNNNNNN**TTACGACCAGTCTAAAAAGCGCC** | Reverse primer for the generation of $Lib_{random}$ |
| p7 | ATATAT<u>GCGGCCGC</u><u>GAAGAC</u>TCGCATNNNNNNNNNNNNNNNNNNNNNNNNN**TTACGACCAGTCTAAAAAGCGCC** | Reverse primer for the generation of 5'-UTR half-library |
| p8 | ATATAT<u>CTGCAG</u><u>GAAGAC</u>CCATGCGNGCNCTNGTNGTNATHCGNCTNTCNCGNGTNACNGAYGCNACN**<u>ACTAGT</u>CCGGAACGTCAGCTGG** | Forward primer for the generation of CDS half-library |
| p9 | **GCCTTTCGTTTTATTTGATGCC** | Reverse primer for the generation of CDS half-library and reverse primer for amplification of 5'-UTR half-library insert |
| p10 | **GGTTATCCAGGCTAAAATCG** | Forward primer for the amplification of 5'-UTR half-library insert |

**Supplementary Table 2: Strains used in this study.**

| Name | Genotype | Description | Source/Reference |
|---|---|---|---|
| *E. coli* TOP10 *ΔrhaA* | *$F^-$ mcrA Δ(mrr-hsdRMS-mcrBC) φ80lacZΔM15 ΔlacX74 nupG recA1 araD139 Δ(ara-leu)7697 galE15 galK16 rpsL($Str^R$) endA1 $\lambda^-$ ΔrhaA* | Rhamnose utilisation-deficient derivative of *E. coli* TOP10. | Hoellerer *et al.* (*1*) |
| *E. coli* TOP10 *ΔrhaA ΔmetZWV* | *$F^-$ mcrA Δ(mrr-hsdRMS-mcrBC) φ80lacZΔM15 ΔlacX74 nupG recA1 araD139 Δ(ara-leu)7697 galE15 galK16 rpsL($Str^R$) endA1 $\lambda^-$ ΔrhaA ΔmetZWV::specR* | Derivative of *E. coli* TOP10 *ΔrhaA* with deleted *metZWV* locus. | This study (**Methods**) |

**Supplementary Table 3: Plasmids used in this study.**

| Name | Description | Source |
|---|---|---|
| pSEVA291 | contains a pBR322 replicon, a kanamycin resistance cassette, and a multiple cloning site | Silva-Rocha *et al.* (*2*) |
| pSEVA361 | Backbone for ptRNA$^{fMet}$ plasmids; contains a p15A replicon, a chloramphenicol resistance cassette, and a multiple cloning site | Silva-Rocha *et al.* (*2*) |
| pASPIre3 | Derivative of pSEVA291; contains a *bxb1-sfGFP* gene (translational fusion) under control of $P_{rha}$ and an *attB-/attP*-flanked 150 bp stretch of silent DNA | Hoellerer *et al.* (*1*) |
| pASPIre4 | Derivative of pASPIre3 with SpeI site in the CDS of *bxb1*. PCR template and backbone for all 5'-UTR and CDS libraries | This study |
| pASPIre4$_{lib}$ | Derivative of pASPIre4 with randomised 5'-UTR and CDS | This study |
| ptRNA$^{fMet-A37}$ | Derivative of pSEVA361 with expression cassette for tRNA$^{fMet}$ with the native A at position 37 | This study |
| ptRNA $^{fMet-A37C}$ | Derivative of ptRNA$^{fMet-A37}$ with A37C substitution | This study |
| ptRNA $^{fMet-A37G}$ | Derivative of ptRNA$^{fMet-A37}$ with A37G substitution | This study |
| ptRNA $^{fMet-A37U}$ | Derivative of ptRNA$^{fMet-A37}$ with A37T/U substitution | This study |

**Supplementary Table 4: DNA adapters for NGS.** Adapters contain NcoI- and SpeI-compatible overhangs (orange) for ligation. Binding regions for Illumina flow cell (blue) and sequencing primers (green) are highlighted. Sample-specific indices are shown in red. P: 5'-phosphorylation.

| Name | Sequence |
|---|---|
| NcoI$_1$ | 5′ AATGATACGGCGACCACCGAGATCTACACTCTTTCCCTACACGACGCTCTTCCGATCTTATCACGC 3′<br>3′ TTACTATGCCGCTGGTGGCTCTAGATGTGAGAAAGGGATGTGCTGCGAGAAGGCTAGAATAGTGCGGTAC-P 5′ |
| NcoI$_2$ | 5′ AATGATACGGCGACCACCGAGATCTACACTCTTTCCCTACACGACGCTCTTCCGATCTATCGATGTC 3′<br>3′ TTACTATGCCGCTGGTGGCTCTAGATGTGAGAAAGGGATGTGCTGCGAGAAGGCTAGATAGCTACAGGTAC-P 5′ |
| NcoI$_3$ | 5′ AATGATACGGCGACCACCGAGATCTACACTCTTTCCCTACACGACGCTCTTCCGATCTGATCTTGTAC 3′<br>3′ TTACTATGCCGCTGGTGGCTCTAGATGTGAGAAAGGGATGTGCTGCGAGAAGGCTAGACTAGAACATGGTAC-P 5′ |
| NcoI$_4$ | 5′ AATGATACGGCGACCACCGAGATCTACACTCTTTCCCTACACGACGCTCTTCCGATCTCGATGCCAATC 3′<br>3′ TTACTATGCCGCTGGTGGCTCTAGATGTGAGAAAGGGATGTGCTGCGAGAAGGCTAGAGCTACGGTTAGGTAC-P 5′ |
| NcoI$_5$ | 5′ AATGATACGGCGACCACCGAGATCTACACTCTTTCCCTACACGACGCTCTTCCGATCTTCGATACAGTGC 3′<br>3′ TTACTATGCCGCTGGTGGCTCTAGATGTGAGAAAGGGATGTGCTGCGAGAAGGCTAGAAGCTATGTCACGGTAC-P 5′ |
| NcoI$_6$ | 5′ AATGATACGGCGACCACCGAGATCTACACTCTTTCCCTACACGACGCTCTTCCGATCTATCGATACTTGAC 3′<br>3′ TTACTATGCCGCTGGTGGCTCTAGATGTGAGAAAGGGATGTGCTGCGAGAAGGCTAGATAGCTATGAACTGGTAC-P 5′ |
| SpeI$_1$ | 5′ P-CTAGAAATAACGTAAGATCGGAAGAGCACACGTCTGAACTCCAGTCACATCTCGTATGCCGTCTTCTGCTTG 3′<br>3′ TTTATTGCATTCTAGCCTTCTCGTGTGCAGACTTGAGGTCAGTGTAGAGCATACGGCAGAAGACGAAC 5′ |
| SpeI$_2$ | 5′ P-CTAGATTCTTGAAATAGATCGGAAGAGCACACGTCTGAACTCCAGTCACATCTCGTATGCCGTCTTCTGCTTG 3′<br>3′ TAAGAACTTTATCTAGCCTTCTCGTGTGCAGACTTGAGGTCAGTGTAGAGCATACGGCAGAAGACGAAC 5′ |
| SpeI$_3$ | 5′ P-CTAGAGGCAGATCATCAGATCGGAAGAGCACACGTCTGAACTCCAGTCACATCTCGTATGCCGTCTTCTGCTTG 3′<br>3′ TCCGTCTAGTAGTCTAGCCTTCTCGTGTGCAGACTTGAGGTCAGTGTAGAGCATACGGCAGAAGACGAAC 5′ |
| SpeI$_4$ | 5′ P-CTAGACTATGTTAATCGAGATCGGAAGAGCACACGTCTGAACTCCAGTCACATCTCGTATGCCGTCTTCTGCTTG 3′<br>3′ TGATACAATTAGCTCTAGCCTTCTCGTGTGCAGACTTGAGGTCAGTGTAGAGCATACGGCAGAAGACGAAC 5′ |
| SpeI$_5$ | 5′ P-CTAGAGTTGACGCATCGAAGATCGGAAGAGCACACGTCTGAACTCCAGTCACATCTCGTATGCCGTCTTCTGCTTG 3′<br>3′ TCAACTGCGTAGCTTCTAGCCTTCTCGTGTGCAGACTTGAGGTCAGTGTAGAGCATACGGCAGAAGACGAAC 5′ |
| SpeI$_6$ | 5′ P-CTAGAATCTACGAATCGATAGATCGGAAGAGCACACGTCTGAACTCCAGTCACATCTCGTATGCCGTCTTCTGCTTG 3′<br>3′ TTAGATGCTTAGCTATCTAGCCTTCTCGTGTGCAGACTTGAGGTCAGTGTAGAGCATACGGCAGAAGACGAAC 5′ |

**Supplementary Table 5: NGS runs and sample-specific adapter combinations from this study.**

| NGS run | | Library | Sample number | Time point (min) | Forward adapter | Reverse adapter |
|---|---|---|---|---|---|---|
| 1 | | $Lib_{random}$ | 1 | 0 | $NcoI_1$ | $SpeI_1$ |
| | | | 2 | 95 | $NcoI_2$ | $SpeI_2$ |
| | | | 3 | 225 | $NcoI_3$ | $SpeI_3$ |
| | | | 4 | 290 | $NcoI_4$ | $SpeI_4$ |
| | | | 5 | 360 | $NcoI_5$ | $SpeI_5$ |
| | | | 6 | 480 | $NcoI_6$ | $SpeI_6$ |
| | | $Lib_{comb1}$ | 1 | 0 | $NcoI_1$ | $SpeI_2$ |
| | | | 2 | 95 | $NcoI_2$ | $SpeI_3$ |
| | | | 3 | 225 | $NcoI_3$ | $SpeI_4$ |
| | | | 4 | 290 | $NcoI_4$ | $SpeI_5$ |
| | | | 5 | 360 | $NcoI_5$ | $SpeI_6$ |
| | | | 6 | 480 | $NcoI_6$ | $SpeI_1$ |
| | | $Lib_{comb2}$ | 1 | 0 | $NcoI_1$ | $SpeI_3$ |
| | | | 2 | 95 | $NcoI_2$ | $SpeI_4$ |
| | | | 3 | 225 | $NcoI_3$ | $SpeI_5$ |
| | | | 4 | 290 | $NcoI_4$ | $SpeI_6$ |
| | | | 5 | 360 | $NcoI_5$ | $SpeI_1$ |
| | | | 6 | 480 | $NcoI_6$ | $SpeI_2$ |
| | | Standard RBSs | 1 | 0 | $NcoI_1$ | $SpeI_4$ |
| | | | 2 | 95 | $NcoI_2$ | $SpeI_5$ |
| | | | 3 | 225 | $NcoI_3$ | $SpeI_6$ |
| | | | 4 | 290 | $NcoI_4$ | $SpeI_1$ |
| | | | 5 | 360 | $NcoI_5$ | $SpeI_2$ |
| | | | 6 | 480 | $NcoI_6$ | $SpeI_3$ |
| 2 | | $Lib_{fact}$ (replicate 1) | 1 | 0 | $NcoI_1$ | $SpeI_1$ |
| | | | 2 | 95 | $NcoI_2$ | $SpeI_2$ |
| | | | 3 | 225 | $NcoI_3$ | $SpeI_3$ |
| | | | 4 | 290 | $NcoI_4$ | $SpeI_4$ |
| | | | 5 | 360 | $NcoI_5$ | $SpeI_5$ |
| | | | 6 | 480 | $NcoI_6$ | $SpeI_6$ |
| | | $Lib_{fact}$ (replicate 2) | 1 | 0 | $NcoI_1$ | $SpeI_6$ |
| | | | 2 | 95 | $NcoI_2$ | $SpeI_5$ |
| | | | 3 | 225 | $NcoI_3$ | $SpeI_4$ |
| | | | 4 | 290 | $NcoI_4$ | $SpeI_3$ |
| | | | 5 | 360 | $NcoI_5$ | $SpeI_2$ |
| | | | 6 | 480 | $NcoI_6$ | $SpeI_1$ |
| 3 | | $Lib_{random}$ in WT strain with $ptRNA^{fMet\text{-}A37}$ | 1 | 0 | $NcoI_1$ | $SpeI_1$ |
| | | | 2 | 95 | $NcoI_2$ | $SpeI_2$ |
| | | | 3 | 225 | $NcoI_3$ | $SpeI_3$ |
| | | | 4 | 290 | $NcoI_4$ | $SpeI_4$ |
| | | | 5 | 360 | $NcoI_5$ | $SpeI_5$ |
| | | | 6 | 480 | $NcoI_6$ | $SpeI_6$ |
| | | $Lib_{random}$ in WT strain with $ptRNA^{fMet\text{-}A37G}$ | 1 | 0 | $NcoI_1$ | $SpeI_2$ |
| | | | 2 | 95 | $NcoI_2$ | $SpeI_3$ |
| | | | 3 | 225 | $NcoI_3$ | $SpeI_4$ |
| | | | 4 | 290 | $NcoI_4$ | $SpeI_5$ |
| | | | 5 | 360 | $NcoI_5$ | $SpeI_6$ |
| | | | 6 | 480 | $NcoI_6$ | $SpeI_1$ |

| | | | | | | |
|---|---|---|---|---|---|---|
| 3 | | Lib$_{random}$ in WT strain with ptRNA$^{fMet-A37U}$ | 1 | 0 | NcoI$_{1}$ | SpeI$_{3}$ |
| | | | 2 | 95 | NcoI$_{2}$ | SpeI$_{4}$ |
| | | | 3 | 225 | NcoI$_{3}$ | SpeI$_{5}$ |
| | | | 4 | 290 | NcoI$_{4}$ | SpeI$_{6}$ |
| | | | 5 | 360 | NcoI$_{5}$ | SpeI$_{1}$ |
| | | | 6 | 480 | NcoI$_{6}$ | SpeI$_{2}$ |
| | | Lib$_{random}$ in *ΔmetZWV* strain with ptRNA$^{fMet-A37}$ | 1 | 0 | NcoI$_{1}$ | SpeI$_{4}$ |
| | | | 2 | 95 | NcoI$_{2}$ | SpeI$_{5}$ |
| | | | 3 | 225 | NcoI$_{3}$ | SpeI$_{6}$ |
| | | | 4 | 290 | NcoI$_{4}$ | SpeI$_{1}$ |
| | | | 5 | 360 | NcoI$_{5}$ | SpeI$_{2}$ |
| | | | 6 | 480 | NcoI$_{6}$ | SpeI$_{3}$ |
| | | Lib$_{random}$ in *ΔmetZWV* strain with ptRNA$^{fMet-A37G}$ | 1 | 0 | NcoI$_{1}$ | SpeI$_{5}$ |
| | | | 2 | 95 | NcoI$_{2}$ | SpeI$_{6}$ |
| | | | 3 | 225 | NcoI$_{3}$ | SpeI$_{1}$ |
| | | | 4 | 290 | NcoI$_{4}$ | SpeI$_{2}$ |
| | | | 5 | 360 | NcoI$_{5}$ | SpeI$_{3}$ |
| | | | 6 | 480 | NcoI$_{6}$ | SpeI$_{4}$ |
| | | Lib$_{random}$ in *ΔmetZWV* strain with ptRNA$^{fMet-A37U}$ | 1 | 0 | NcoI$_{1}$ | SpeI$_{6}$ |
| | | | 2 | 95 | NcoI$_{2}$ | SpeI$_{1}$ |
| | | | 3 | 225 | NcoI$_{3}$ | SpeI$_{2}$ |
| | | | 4 | 290 | NcoI$_{4}$ | SpeI$_{3}$ |
| | | | 5 | 360 | NcoI$_{5}$ | SpeI$_{4}$ |
| | | | 6 | 480 | NcoI$_{6}$ | SpeI$_{5}$ |

**Supplementary Table 6: Codon weight and frequencies for calculation of CAI and tAI.** w(CAI)/w(tAI): weight for the calculation of CAI (Sharp *et al.* (*3*))/tAI (dos Reis *et al.* (*4*)). *E. coli* fraction: fraction of each triplet amongst all triplets coding for the same amino acid in the *E. coli* genome (according to: https://www.kazusa.or.jp/codon/).

| **Triplet** | **Amino acid** | **w(CAI)** | **w(tAI)** | **Rel. codon frequency** |
|---|---|---|---|---|
| GCA | Ala | 0.586 | 0.375 | 0.270 |
| GCC | Ala | 0.122 | 0.250 | 0.260 |
| GCG | Ala | 0.424 | 0.120 | 0.250 |
| GCU | Ala | 1.000 | 0.110 | 0.220 |
| AGA | Arg | 0.004 | 0.125 | 0.130 |
| AGG | Arg | 0.002 | 0.165 | 0.070 |
| CGA | Arg | 0.004 | 0.000 | 0.090 |
| CGC | Arg | 0.356 | 0.360 | 0.260 |
| CGG | Arg | 0.004 | 0.125 | 0.150 |
| CGU | Arg | 1.000 | 0.500 | 0.300 |
| AAC | Asn | 1.000 | 0.500 | 0.410 |
| AAU | Asn | 0.051 | 0.220 | 0.590 |
| GAC | Asp | 1.000 | 0.375 | 0.350 |
| GAU | Asp | 0.434 | 0.165 | 0.650 |
| UGC | Cys | 1.000 | 0.125 | 0.480 |
| UGU | Cys | 0.500 | 0.055 | 0.520 |
| CAA | Gln | 0.124 | 0.250 | 0.350 |
| CAG | Gln | 1.000 | 0.330 | 0.650 |

| | | | | |
|---|---|---|---|---|
| GAA | Glu | 1.000 | 0.500 | 0.640 |
| GAG | Glu | 0.259 | 0.160 | 0.360 |
| GGA | Gly | 0.010 | 0.125 | 0.190 |
| GGC | Gly | 0.724 | 0.500 | 0.290 |
| GGG | Gly | 0.019 | 0.165 | 0.180 |
| GGU | Gly | 1.000 | 0.220 | 0.340 |
| CAC | His | 1.000 | 0.125 | 0.370 |
| CAU | His | 0.291 | 0.055 | 0.630 |
| AUA | Ile | 0.003 | 0.163 | 0.210 |
| AUC | Ile | 1.000 | 0.375 | 0.310 |
| AUU | Ile | 0.185 | 0.165 | 0.470 |
| CUA | Leu | 0.007 | 0.125 | 0.060 |
| CUC | Leu | 0.037 | 0.125 | 0.100 |
| CUG | Leu | 1.000 | 0.540 | 0.380 |
| CUU | Leu | 0.042 | 0.055 | 0.150 |
| UUA | Leu | 0.020 | 0.125 | 0.180 |
| UUG | Leu | 0.020 | 0.165 | 0.130 |
| AAA | Lys | 1.000 | 0.750 | 0.710 |
| AAG | Lys | 0.253 | 0.240 | 0.290 |
| AUG | Met | 1.000 | 1.000 | 1.000 |
| UUC | Phe | 1.000 | 0.250 | 0.360 |
| UUU | Phe | 0.296 | 0.110 | 0.640 |
| CCA | Pro | 0.135 | 0.125 | 0.230 |
| CCC | Pro | 0.012 | 0.125 | 0.160 |
| CCG | Pro | 1.000 | 0.165 | 0.370 |
| CCU | Pro | 0.070 | 0.055 | 0.240 |
| AGC | Ser | 0.410 | 0.125 | 0.200 |
| AGU | Ser | 0.085 | 0.055 | 0.180 |
| UCA | Ser | 0.077 | 0.125 | 0.180 |
| UCC | Ser | 0.744 | 0.250 | 0.140 |
| UCG | Ser | 0.017 | 0.165 | 0.110 |
| UCU | Ser | 1.000 | 0.110 | 0.180 |
| ACA | Thr | 0.076 | 0.125 | 0.250 |
| ACC | Thr | 1.000 | 0.250 | 0.310 |
| ACG | Thr | 0.099 | 0.290 | 0.220 |
| ACU | Thr | 0.965 | 0.110 | 0.220 |
| UGG | Trp | 1.000 | 0.165 | 1.000 |
| UAC | Tyr | 1.000 | 0.375 | 0.350 |
| UAU | Tyr | 0.239 | 0.165 | 0.650 |
| GUA | Val | 0.495 | 0.625 | 0.190 |
| GUC | Val | 0.066 | 0.250 | 0.190 |
| GUG | Val | 0.221 | 0.200 | 0.290 |
| GUU | Val | 1.000 | 0.110 | 0.320 |

**Supplementary Table 7: Optimisation of hybridisation window.** Correlation between rTR and the mean hybridisation energy of different sequence windows with different lengths. The mean of the hybridisation energy at position -11 and position -10 (rank 1) was denoted $hyb_{opt}$. The data was ranked by the sum of R and ρ.

| Rank | Window length (nt) | Window start (5'-UTR pos.) | Pearson's R | Spearman's ρ |
|---|---|---|---|---|
| 1 | 2 | -11 | -0.2979 | -0.1696 |
| 2 | 3 | -11 | -0.2979 | -0.1671 |
| 3 | 4 | -12 | -0.2988 | -0.1660 |
| 4 | 3 | -12 | -0.2935 | -0.1652 |
| 5 | 5 | -12 | -0.2953 | -0.1613 |
| 6 | 5 | -13 | -0.2916 | -0.1601 |
| 7 | 6 | -13 | -0.2930 | -0.1586 |
| 8 | 2 | -10 | -0.2884 | -0.1619 |
| 9 | 4 | -11 | -0.2893 | -0.1588 |
| 10 | 7 | -13 | -0.2866 | -0.1524 |
| 11 | 2 | -12 | -0.2797 | -0.1585 |
| 12 | 4 | -13 | -0.2814 | -0.1557 |
| 13 | 6 | -12 | -0.2845 | -0.1521 |
| 14 | 7 | -14 | -0.2833 | -0.1513 |
| 15 | 8 | -14 | -0.2810 | -0.1479 |
| 16 | 6 | -14 | -0.2776 | -0.1497 |
| 17 | 3 | -10 | -0.2747 | -0.1507 |
| 18 | 5 | -11 | -0.2739 | -0.1467 |
| 19 | 8 | -13 | -0.2741 | -0.1425 |
| 20 | 9 | -14 | -0.2720 | -0.1403 |

### Supplementary Notes

**Supplementary Note 1: Sequence of spectinomycin resistance gene (*specR*) including constitutive promoter used for knockout of *metZWV*.** The start and stop codon of *specR* are underlined.

TGATCGGCACGTAAGAGGTTCCAACTTTCACCATAATGAAATAAGATCACTACCGGGCGTATTTTTTGAGTTATC
GAGATTTTCAGGAGCTAAGGAAGCTACATATGAGTGAAAAAGTGCCCGCCGAGATTTCGGTGCAACTATCACAAG
CACTCAACGTCATCGGGCGCCACTTGGAGTCGACGTTGCTGGCCGTGCATTTGTACGGCTCCGCACTGGATGGCG
GATTGAAACCGTACAGTGATATTGATTTGCTGGTGACTGTAGCTGCACCGCTCAATGATGCCGTGCGGCAAGCCC
TGCTCGTCGATCTCTTGGAGGTTTCAGCTTCCCCTGGCCAAAACAAGGCACTCCGCGCCTTGGAAGTGACCATCG
TCGTGCACAGTGACATCGTACCTTGGCGTTATCCGGCCAGGCGGGAACTGCAGTTCGGAGAGTGGCAGCGCAAAG
ACATCCTTGCGGGCATCTTCGAGCCCGCCACAACCGATTCTGACTTGGCGATTCTGCTAACAAAGGCAAAGCAAC
ATAGCGTCGTCTTGGCAGGTTCAGCAGCGAAGGATCTCTTCAGCTCAGTCCCAGAAAGCGATCTATTCAAGGCAC
TGGCCGATACTCTGAAGCTATGGAACTCGCCGCCAGATTGGGCGGGCGATGAGCGGAATGTAGTGCTTACTTTGT
CTCGTATCTGGTACACCGCAGCAACCGGCAAGATCGCGCCAAAGGATGTTGCTGCCACTTGGGCAATGGCACGCT
TGCCAGCTCAACATCAGCCCATCCTGTTGAATGCCAAGCGGGCTTATCTTGGGCAAGAAGAAGATTATTTGCCCG
CTCGTGCGGATCAGGTGGCGGCGCTCATTAAATTCGTGAAGTATGAAGCAGTTAAACTGCTTGGTGCCAGCCAAT
AA

**Supplementary Note 2: Sequence of pASPIre4.** The displayed sequence corresponds to the insert between HindIII and PacI restriction sites (underlined) in pSEVA291. A graphical representation of pASPIre4 is shown in **Supplementary Figure 1**.

AAGCTTCACATCTGCAGTAATCGGCCGGCTTGTCGACGACGGCGGTCTCCGTCGTCAGGATCATCCGGGCATCGC
TTAGTCACCTTTGGGCCACGGTCCGCTACCTTACAGGAATAGTACTCGTCCTTTAATTTGGAATGAACCATGGCA
GTCAGTTGTGTTGCGTTTCTTCGACCTAGTACTCGCTCCCTTAGGAGAAAGACAGATAGCTTCTTACCCGGGGTT
TGTACCGTACACCACTGAGACCGCGGTGGTTGACCAGACAAACCACGAAGGTTCTGTTAAGTAACTGAACCCAAT
GTCGTTAGTGACGCTTACCTCTTAAGAGGTCACTGACCTAACAGGATCCCACCACAATTCAGCAAATTGTGAACA
TCATCACGTTCATCTTTCCCTGGTTGCCAATGGCCCATTTTCCTGTCAGTAACGAGAAGGTCGCGAATTCAGGCG
CTTTTTAGACTGGTCGTAATGAAGAGCTCAATAAATATTTAATTTATCTCAGAAAGGCTAAGACATGCGAGCACT
GGTTGTTATTCGTCTGAGCCGTGTTACCGATGCAACCACTAGTCCGGAACGTCAGCTGGAAAGCTGTCAGCAGCT
GTGTGCACAGCGTGGTTGGGATGTTGTTGGTGTTGCAGAGGATCTGGATGTTAGCGGTGCAGTTGATCCGTTTGA
TCGTAAACGTCGTCCGAATCTGGCACGTTGGCTGGCATTTGAAGAACAGCCGTTTGATGTTATTGTTGCCTATCG
TGTTGATCGTCTGACCCGTAGCATTCGTCATCTGCAACAGCTGGTTCATTGGGCAGAAGATCATAAAAAACTGGT
TGTGAGCGCAACCGAAGCACATTTTGATACCACCACCCCGTTTGCAGCAGTTGTTATTGCACTGATGGGCACCGT
TGCACAGATGGAACTGGAAGCAATTAAAGAACGTAATCGTAGCGCAGCCCATTTTAACATTCGTGCAGGTAAATA
TCGTGGTAGCCTGCCTCCGTGGGGTTATCTGCCGACGCGTGTAGATGGTGAATGGCGTCTGGTTCCTGACCCGGT
TCAGCGTGAACGTATTCTGGAAGTATATCATCGTGTGGTGGATAATCATGAACCGCTGCATCTGGTTGCACATGA
TCTGAATCGTCGTGGTGTTCTGAGTCCCAAAGATTATTTTGCTCAGCTGCAAGGTCGTGAACCGCAGGGTCGTGA
ATGGTCTGCAACCGCACTGAAACGTAGCATGATTAGCGAAGCAATGCTGGGTTATGCAACCCTGAATGGTAAAAC
CGTTCGTGATGATGATGGTGCACCGCTGGTTCGTGCAGAACCGATTCTGACACGTGAACAGCTGGAAGCACTGCG
TGCCGAACTGGTTAAAACCAGCCGTGCAAAACCGGCAGTTAGCACCCCGAGCCTGCTGCTGCGTGTTCTGTTTTG
TGCAGTTTGTGGTGAACCGGCATACAAATTTGCCGGTGGTGGTCGTAAACATCCGCGTTATCGTTGTCGTAGCAT
GGGTTTTCCGAAACATTGTGGTAATGGTACAGTTGCAATGGCAGAATGGGATGCATTTTGCGAAGAACAGGTTCT
GGATCTGCTGGGTGATGCCGAACGTCTGGAAAAAGTTTGGGTTGCAGGTAGCGATAGCGCAGTTGAACTGGCCGA
AGTTAATGCGGAACTGGTCGATCTCACCAGTCTGATTGGAAGTCCCGCATATCGTGCGGGTAGTCCTCAGCGTGA
AGCACTGGATGCACGTATTGCAGCACTGGCAGCACGTCAAGAAGAACTGGAAGGTCTGGAAGCACGTCCGAGCGG
TTGGGAATGGCGTGAAACAGGTCAGCGTTTTGGTGATTGGTGGCGTGAGCAGGATACCGCAGCAAAAAATACCTG
GCTGCGTAGTATGAATGTTCGCCTGACCTTTGATGTTCGCGGTGGCCTGACCCGCACCATTGATTTTGGCGATCT
GCAAGAATATGAACAGCATCTGCGTCTGGGTAGCGTTGTTGAACGTCTGCATACCGGCATGAGCACCGGCGGTGG
CAGCGGCGGTTCTGGTGGCTCTAGCAAAGGAGAAGAACTTTTCACTGGAGTTGTCCCAATTCTTGTTGAATTAGA
TGGTGATGTTAATGGGCACAAATTTTCTGTCCGTGGAGAGGGTGAAGGTGATGCTACAAACGGAAAACTCACCCT

TAAATTTATTTGCACTACTGGAAAACTACCTGTTCCGTGGCCAACACTTGTCACTACTCTGACCTATGGTGTTCA
ATGCTTTTCCCGTTATCCGGATCACATGAAACGGCATGACTTTTTCAAGAGTGCCATGCCCGAAGGTTATGTACA
GGAACGCACTATATCTTTCAAAGATGACGGGACCTACAAGACGCGTGCTGAAGTCAAGTTTGAAGGTGATACCCT
TGTTAATCGTATCGAGTTAAAGGGTATTGATTTTAAAGAAGATGGAAACATTCTTGGACACAAACTCGAGTACAA
CTTTAACTCACACAATGTATACATCACGGCAGACAAACAAAAGAATGGAATCAAAGCTAACTTCAAAATTCGCCA
CAACGTTGAAGATGGTTCCGTTCAACTAGCAGACCATTATCAACAAAATACTCCAATTGGCGATGGCCCTGTCCT
TTTACCAGACAACCATTACCTGTCGACACAATCTGTCCTTTCGAAAGATCCCAACGAAAAGCGTGACCACATGGT
CCTTCTTGAGTTTGTAACTGCTGCTGGGATTACACATGGCATGGATGAACTCTACAAAAGGCCTGCTGCTAACGA
CGAAAACTACGCTCTGGCTGCTTAATAAGCGGCCGCGGCCTAGGCGGCCTCCTGTGTGAAATTGTTATCCGCTTT
AATTAA

**Supplementary Note 3: Sequence of pASPIre4$_{lib}$.** The displayed sequence corresponds to the insert between PstI and NotI restriction sites (underlined) in pASPIre4. A graphical representation of pASPIre4$_{lib}$ is shown in **Supplementary Figure 2**.

CTGCAGTAATCGGCCGGCTTGTCGACGACGGCGGTCTCCGTCGTCAGGATCATCCGGGCATCGCTTAGTCACCTT
TGGGCCACGGTCCGCTACCTTACAGGAATAGTACTCGTCCTTTAATTTGGAATGAACCATGGCAGTCAGTTGTGT
TGCGTTTCTTCGACCTAGTACTCGCTCCCTTAGGAGAAAGACAGATAGCTTCTTACCCGGGGTTTGTACCGTACA
CCACTGAGACCGCGGTGGTTGACCAGACAAACCACGAAGGTTCTGTTAAGTAACTGAACCCAATGTCGTTAGTGA
CGCTTACCTCTTAAGAGGTCACTGACCTAACAGGATCCCACCACAATTCAGCAAATTGTGAACATCATCACGTTC
ATCTTTCCCTGGTTGCCAATGGCCCATTTTCCTGTCAGTAACGAGAAGGTCGCGAATTCAGGCGCTTTTTAGACT
GGTCGTAANNNNNNNNNNNNNNNNNNNNNNNNNATGCGNGCNCTNGTNGTNATHCGNCTNTCNCGNGTNACNGAY
GCNACNACTAGTCCGGAACGTCAGCTGGAAAGCTGTCAGCAGCTGTGTGCACAGCGTGGTTGGGATGTTGTTGGT
GTTGCAGAGGATCTGGATGTTAGCGGTGCAGTTGATCCGTTTGATCGTAAACGTCGTCCGAATCTGGCACGTTGG
CTGGCATTTGAAGAACAGCCGTTTGATGTTATTGTTGCCTATCGTGTTGATCGTCTGACCCGTAGCATTCGTCAT
CTGCAACAGCTGGTTCATTGGGCAGAAGATCATAAAAAACTGGTTGTGAGCGCAACCGAAGCACATTTTGATACC
ACCACCCCGTTTGCAGCAGTTGTTATTGCACTGATGGGCACCGTTGCACAGATGGAACTGGAAGCAATTAAAGAA
CGTAATCGTAGCGCAGCCCATTTTAACATTCGTGCAGGTAAATATCGTGGTAGCCTGCCTCCGTGGGGTTATCTG
CCGACGCGTGTAGATGGTGAATGGCGTCTGGTTCCTGACCCGGTTCAGCGTGAACGTATTCTGGAAGTATATCAT
CGTGTGGTGGATAATCATGAACCGCTGCATCTGGTTGCACATGATCTGAATCGTCGTGGTGTTCTGAGTCCCAAA
GATTATTTTGCTCAGCTGCAAGGTCGTGAACCGCAGGGTCGTGAATGGTCTGCAACCGCACTGAAACGTAGCATG
ATTAGCGAAGCAATGCTGGGTTATGCAACCCTGAATGGTAAAACCGTTCGTGATGATGATGGTGCACCGCTGGTT
CGTGCAGAACCGATTCTGACACGTGAACAGCTGGAAGCACTGCGTGCCGAACTGGTTAAAACCAGCCGTGCAAAA
CCGGCAGTTAGCACCCCGAGCCTGCTGCTGCGTGTTCTGTTTTGTGCAGTTTGTGGTGAACCGGCATACAAATTT
GCCGGTGGTGGTCGTAAACATCCGCGTTATCGTTGTCGTAGCATGGGTTTTCCGAAACATTGTGGTAATGGTACA
GTTGCAATGGCAGAATGGGATGCATTTTGCGAAGAACAGGTTCTGGATCTGCTGGGTGATGCCGAACGTCTGGAA
AAAGTTTGGGTTGCAGGTAGCGATAGCGCAGTTGAACTGGCCGAAGTTAATGCGGAACTGGTCGATCTCACCAGT
CTGATTGGAAGTCCCGCATATCGTGCGGGTAGTCCTCAGCGTGAAGCACTGGATGCACGTATTGCAGCACTGGCA
GCACGTCAAGAAGAACTGGAAGGTCTGGAAGCACGTCCGAGCGGTTGGGAATGGCGTGAAACAGGTCAGCGTTTT
GGTGATTGGTGGCGTGAGCAGGATACCGCAGCAAAAAATACCTGGCTGCGTAGTATGAATGTTCGCCTGACCTTT
GATGTTCGCGGTGGCCTGACCCGCACCATTGATTTTGGCGATCTGCAAGAATATGAACAGCATCTGCGTCTGGGT
AGCGTTGTTGAACGTCTGCATACCGGCATGAGCACCGGCGGTGGCAGCGGCGGTTCTGGTGGCTCTAGCAAAGGA
GAAGAACTTTTCACTGGAGTTGTCCCAATTCTTGTTGAATTAGATGGTGATGTTAATGGGCACAAATTTTCTGTC
CGTGGAGAGGGTGAAGGTGATGCTACAAACGGAAAACTCACCCTTAAATTTATTTGCACTACTGGAAAACTACCT
GTTCCGTGGCCAACACTTGTCACTACTCTGACCTATGGTGTTCAATGCTTTTCCCGTTATCCGGATCACATGAAA
CGGCATGACTTTTTCAAGAGTGCCATGCCCGAAGGTTATGTACAGGAACGCACTATATCTTTCAAAGATGACGGG
ACCTACAAGACGCGTGCTGAAGTCAAGTTTGAAGGTGATACCCTTGTTAATCGTATCGAGTTAAAGGGTATTGAT
TTTAAAGAAGATGGAAACATTCTTGGACACAAACTCGAGTACAACTTTAACTCACACAATGTATACATCACGGCA
GACAAACAAAAGAATGGAATCAAAGCTAACTTCAAAATTCGCCACAACGTTGAAGATGGTTCCGTTCAACTAGCA
GACCATTATCAACAAAATACTCCAATTGGCGATGGCCCTGTCCTTTTACCAGACAACCATTACCTGTCGACACAA
TCTGTCCTTTCGAAAGATCCCAACGAAAAGCGTGACCACATGGTCCTTCTTGAGTTTGTAACTGCTGCTGGGATT
ACACATGGCATGGATGAACTCTACAAAAGGCCTGCTGCTAACGACGAAAACTACGCTCTGGCTGCTTAATAAGCG
GCCGC

**Supplementary Note 4: Sequence of 5'-UTR half-library.** The displayed sequence corresponds to the insert between PstI and NotI restriction sites (underlined) in pASPIre4. The BbsI restriction site for scarless cloning of 5'-UTR-CDS combinations is marked in orange. A graphical representation is available in **Supplementary Figure 4**.

CTGCAGTAATCGGCCGGCTTGTCGACGACGGCGGTCTCCGTCGTCAGGATCATCCGGGCATCGCTTAGTCACCTT
TGGGCCACGGTCCGCTACCTTACAGGAATAGTACTCGTCCTTTAATTTGGAATGAACCATGGCAGTCAGTTGTGT
TGCGTTTCTTCGACCTAGTACTCGCTCCCTTAGGAGAAAGACAGATAGCTTCTTACCCGGGGTTTGTACCGTACA
CCACTGAGACCGCGGTGGTTGACCAGACAAACCACGAAGGTTCTGTTAAGTAACTGAACCCAATGTCGTTAGTGA
CGCTTACCTCTTAAGAGGTCACTGACCTAACAGGATCCCACCACAATTCAGCAAATTGTGAACATCATCACGTTC
ATCTTTCCCTGGTTGCCAATGGCCCATTTTCCTGTCAGTAACGAGAAGGTCGCGAATTCAGGCGCTTTTTAGACT
GGTCGTAANNNNNNNNNNNNNNNNNNNNNNNNNATGCGAGTCTTCGCGGCCGC

**Supplementary Note 5: Sequence of CDS half-library.** The displayed sequence corresponds to the insert between PstI and NotI restriction sites (underlined) in pASPIre4. The BbsI restriction site for scarless cloning of 5'-UTR-CDS combinations is marked in orange. A graphical representation is available in **Supplementary Figure 5**.

CTGCAGGAAGACCCATGCGNGCNCTNGTNGTNATHCGNCTNTCNCGNGTNACNGAYGCNACNACTAGTCCGGAAC
GTCAGCTGGAAAGCTGTCAGCAGCTGTGTGCACAGCGTGGTTGGGATGTTGTTGGTGTTGCAGAGGATCTGGATG
TTAGCGGTGCAGTTGATCCGTTTGATCGTAAACGTCGTCCGAATCTGGCACGTTGGCTGGCATTTGAAGAACAGC
CGTTTGATGTTATTGTTGCCTATCGTGTTGATCGTCTGACCCGTAGCATTCGTCATCTGCAACAGCTGGTTCATT
GGGCAGAAGATCATAAAAAACTGGTTGTGAGCGCAACCGAAGCACATTTTGATACCACCACCCCGTTTGCAGCAG
TTGTTATTGCACTGATGGGCACCGTTGCACAGATGGAACTGGAAGCAATTAAAGAACGTAATCGTAGCGCAGCCC
ATTTTAACATTCGTGCAGGTAAATATCGTGGTAGCCTGCCTCCGTGGGGTTATCTGCCGACGCGTGTAGATGGTG
AATGGCGTCTGGTTCCTGACCCGGTTCAGCGTGAACGTATTCTGGAAGTATATCATCGTGTGGTGGATAATCATG
AACCGCTGCATCTGGTTGCACATGATCTGAATCGTCGTGGTGTTCTGAGTCCCAAAGATTATTTTGCTCAGCTGC
AAGGTCGTGAACCGCAGGGTCGTGAATGGTCTGCAACCGCACTGAAACGTAGCATGATTAGCGAAGCAATGCTGG
GTTATGCAACCCTGAATGGTAAAACCGTTCGTGATGATGATGGTGCACCGCTGGTTCGTGCAGAACCGATTCTGA
CACGTGAACAGCTGGAAGCACTGCGTGCCGAACTGGTTAAAACCAGCCGTGCAAAACCGGCAGTTAGCACCCCGA
GCCTGCTGCTGCGTGTTCTGTTTTGTGCAGTTTGTGGTGAACCGGCATACAAATTTGCCGGTGGTGGTCGTAAAC
ATCCGCGTTATCGTTGTCGTAGCATGGGTTTTCCGAAACATTGTGGTAATGGTACAGTTGCAATGGCAGAATGGG
ATGCATTTTGCGAAGAACAGGTTCTGGATCTGCTGGGTGATGCCGAACGTCTGGAAAAAGTTTGGGTTGCAGGTA
GCGATAGCGCAGTTGAACTGGCCGAAGTTAATGCGGAACTGGTCGATCTCACCAGTCTGATTGGAAGTCCCGCAT
ATCGTGCGGGTAGTCCTCAGCGTGAAGCACTGGATGCACGTATTGCAGCACTGGCAGCACGTCAAGAAGAACTGG
AAGGTCTGGAAGCACGTCCGAGCGGTTGGGAATGGCGTGAAACAGGTCAGCGTTTTGGTGATTGGTGGCGTGAGC
AGGATACCGCAGCAAAAAATACCTGGCTGCGTAGTATGAATGTTCGCCTGACCTTTGATGTTCGCGGTGGCCTGA
CCCGCACCATTGATTTTGGCGATCTGCAAGAATATGAACAGCATCTGCGTCTGGGTAGCGTTGTTGAACGTCTGC
ATACCGGCATGAGCACCGGCGGTGGCAGCGGCGGTTCTGGTGGCTCTAGCAAAGGAGAAGAACTTTTCACTGGAG
TTGTCCCAATTCTTGTTGAATTAGATGGTGATGTTAATGGGCACAAATTTTCTGTCCGTGGAGAGGGTGAAGGTG
ATGCTACAAACGGAAAACTCACCCTTAAATTTATTTGCACTACTGGAAAACTACCTGTTCCGTGGCCAACACTTG
TCACTACTCTGACCTATGGTGTTCAATGCTTTTCCCGTTATCCGGATCACATGAAACGGCATGACTTTTTCAAGA
GTGCCATGCCCGAAGGTTATGTACAGGAACGCACTATATCTTTCAAAGATGACGGGACCTACAAGACGCGTGCTG
AAGTCAAGTTTGAAGGTGATACCCTTGTTAATCGTATCGAGTTAAAGGGTATTGATTTTAAAGAAGATGGAAACA
TTCTTGGACACAAACTCGAGTACAACTTTAACTCACACAATGTATACATCACGGCAGACAAACAAAAGAATGGAA
TCAAAGCTAACTTCAAAATTCGCCACAACGTTGAAGATGGTTCCGTTCAACTAGCAGACCATTATCAACAAAATA
CTCCAATTGGCGATGGCCCTGTCCTTTTACCAGACAACCATTACCTGTCGACACAATCTGTCCTTTCGAAAGATC
CCAACGAAAAGCGTGACCACATGGTCCTTCTTGAGTTTGTAACTGCTGCTGGGATTACACATGGCATGGATGAAC
TCTACAAAAGGCCTGCTGCTAACGACGAAAACTACGCTCTGGCTGCTTAATAAGCGGCCGC

**Supplementary Note 6: Sequence of ptRNA$^{fMet}$ variants.** The displayed sequence corresponds to the insert between KpnI and SpeI restriction sites (underlined) in pSEVA361. The mutated position 37 in tRNA$^{fMet}$ is highlighted in bold. A graphical representation is available in **Supplementary Figure 6**.

GGTACCAAAATAACACCCTGCTTAATTAAAGCGGATAACAATTTCACACAGGAGGCCGCCTAGGCCGCGGCCGCG
CGAATTCGAGCTCGGTACCAAAATAACACCCTGCTTAATTAACTGATGATGAGCCTGGATTTCCGCTCTCACTGA
ATTTTTATGCAAAATAAATGAGTTTTCATTTAATCATCTTTTATCGGAGACAGGAAGAGTTTAGTGTGTTTTTTG
TAAAATAATGCGCTTAAGGGAGAGCAGGAGAAGGCAAAAGTATTCAACAAATGAAAGTGAACTGGATATTCATTC
ACATGATTAGCAATAAACGTTGACAAAATGTGGCGTGGATCACTATAATGCCTGCAGATTTTACGTCCCGTCTCG
GTACACCAAATCCCAGCAGTATTTGCATTTTTTACCCAAAACGAGTAGAATTTGCCACGTTTCAGGCGCGGGGTG
GAGCAGCCTGGTAGCTCGTCGGGCTCAT**N**ACCCGAAGGTCGTCGGTTCAAATCCGGCCCCCGCAACCACTTTCCC
TTAGAGTCCTTTTTCAAATATACTGTGAAGACTTCGGCCTTCGTAGTGGGATTTGAAAAAATCCTTCTGGAAAGT
GCTCCAGACCGCAGTTGCGGTTATAGGGTTCAGTTATATAAAGCCCCGATTTATCGGGGTTTTTTGTTATCTGAC
TACAGAATAACTGGGCTTTAGGCCCTTTTTTTATGTCTTGGGGGTGGGCACTAGT

### References (Supplementary Information)


1. S. Hollerer *et al.*, Large-scale DNA-based phenotypic recording and deep learning enable highly accurate sequence-function mapping. *Nat Commun* **11**, 3551 (2020).
2. R. Silva-Rocha *et al.*, The Standard European Vector Architecture (SEVA): a coherent platform for the analysis and deployment of complex prokaryotic phenotypes. *Nucleic Acids Res* **41**, D666-675 (2013).
3. P. M. Sharp, W. H. Li, The codon Adaptation Index--a measure of directional synonymous codon usage bias, and its potential applications. *Nucleic Acids Res* **15**, 1281-1295 (1987).
4. M. dos Reis, L. Wernisch, R. Savva, Unexpected correlations between gene expression and codon usage bias from microarray data for the whole Escherichia coli K-12 genome. *Nucleic Acids Res* **31**, 6976-6985 (2003).
